# Structural molecular and developmental evidence for cell-type diversity in cnidarian mechanosensory neurons

**DOI:** 10.1101/2024.11.20.624413

**Authors:** Julia Baranyk, Kristen Malir, Miguel A. P. Silva, Sakura Rieck, Gracie Scheve, Nagayasu Nakanishi

## Abstract

Deploying a conserved mechanosensory neuron known as the concentric hair cell, cnidarians have evolved diverse mechanoreceptors from hydroid filiform tentacles to jellyfish statocysts. However, it is unknown whether cnidarian mechanoreceptor evolution has relied solely on repurposing a single ancestral mechanosensory neuron type. Here we report evidence for cell-type diversity of mechanosensory neurons in sea-anemone cnidarian *Nematostella vectensis.* Uncovered in the ectoderm of feeding tentacles are conventional type I hair cells and previously unrecognized type II hair cells differing in the structure of apical sensory apparatus and synapses. Moreover, we identify TRP channel-encoding gene *polycystin-1* as a type-II-hair-cell-specific essential mediator of gentle touch response. Ontogenically, type I and type II hair cells derive from distinct postmitotic precursors that begin forming at different phases of larval development. Taken together, our findings suggest that anatomically, molecularly, and developmentally distinct mechanosensory neurons diversified within Cnidaria, or prior to the divergence of Cnidaria and Bilateria.

## Introduction

United by the presence of a stinging cell type known as the cnidocyte, Cnidaria is an early-branching animal group represented by sea anemones, corals and jellyfishes. Cnidaria split from its sister group Bilateria (e.g. chordates and arthropods) in the late Precambrian ^1^ and further diverged into two major lineages with distinct life cycles: Anthozoa and Medusozoa. Anthozoa consists of Octocorallia (sea pens and soft corals) and Hexacorallia (sea anemones and scleractinian corals), while Medusozoa comprises Staurozoa (stalked jellyfishes), Hydrozoa (hydromedusae, hydroids and siphonophores), Scyphozoa (true jellyfish), and Cubozoa (box jellyfish) ^2,3^. Cnidarian embryogenesis typically generates a free-swimming planula larva that transforms into a sessile polyp with a single oral opening surrounded by feeding tentacles.

Polyps proceed to sexual maturity in Anthozoa, while in Medusozoa they undergo another round of transformation to produce free-swimming medusae that are equipped with feeding tentacles and become sexually mature adults. The cnidocyte-loaded tentacles of polyps and medusae function as sensory structures, detecting water vibration and chemicals emanating from nearby prey to coordinate its capture. In addition, some hydroids develop specialized mechanosensory tentacles known as the filiform tentacles that are thought to mediate a bending reaction of the animal in response to mechanical stimulation of the tentacle ^4^, while a variety of medusae form gravity sensors (e.g. statocysts) to control the balance against gravity ^5–7^. Although it is evident that cnidarians evolved diverse mechanosensory structures in parallel with Bilateria, little is known about the cell-type diversity of cnidarian mechanosensory neurons.

In cnidarian mechanosensory structures such as tentacles, the primary sensory neuron thought to transduce mechanical stimuli (e.g. water vibration or touch) is the concentric hair cell that resides in the ectodermal epithelium. The concentric hair cell is characterized by an apical sensory apparatus consisting of a single cilium surrounded by one or multiple collars of stereovilli/microvilli ^8,9^; stereovilli (or stereocilia) are distinguished from microvilli based on the presence of actin rootlets. The concentric hair cell extends basal neuronal processes that likely facilitate the transmission of mechanosensory information to other cells/tissues such as cnidocytes ^8^ and nerve rings of hydrozoan jellyfish (e.g. ^5,6,9,10^), possibly through unusual synapses with large synaptic vesicles (160-1100 nm in diameter) ^11^. Structural evidence for conventional chemical synapses for signal transmission from concentric hair cells to other cells is currently lacking. Nevertheless, consistent with the cells’ purported mechanosensory function, it has been reported that mechanical stimulation of cilia of concentric hair cells, or of the structure that includes the cilia, elicits coordinated movement of hair-cell-bearing tentacles towards the source of the stimulus in hydrozoan and cubozoan polyps as well as in hydrozoan medusae ^9,12,13^, and triggers electrical responses in the nerve ring of hydrozoan medusae ^9^, cnidocytes of hydroid tentacles ^8^, and hair bundle mechanoreceptors – concentric hair cell-support cell complexes – of sea anemone tentacles ^14,15^.

Although cell type diversity of non-bilaterians such as cnidarians is generally believed to be limited relative to bilaterians (e.g. ^16^), morphology of cnidarian hair cells can vary at the subcellular level within a given taxon, raising the possibility that cnidarians have evolved more than one mechanosensory neural cell type. Indeed, the polyp tentacle of the hydrozoan *Sarsia tubulosa* (formerly *Coryne tubulosa*) houses two concentric hair cell types that differ in cilia length (15 µm vs. 1-2 µm) and another concentric hair cell-like cell type with a recessed cilium-microvilli complex ^17^. Likewise, the polyp of box jellyfish *Carybdea marsupialis* has one concentric hair cell type with a collar of 7-9 pronounced stereovilli, and another cell type with a long cilium (40 µm in length) surrounded by rings of short microvilli ^12^. However, these morphological data are also consistent with alternative interpretations that cells having different morphologies of apical mechanosensor represent morphological variants, or different developmental states, of the same cell type. Thus, it remains unclear whether cnidarians have different mechanosensory neural cell types with distinct mechanotransduction machineries and developmental histories, akin to bilaterian conditions (e.g. gentle touch and harsh touch sensory neurons of *C. elegans* worms ^18,19^). Consequently, it is unknown whether cnidarian mechanoreceptor evolution has involved diversification of molecularly and developmentally distinct mechanosensory neurons, or been solely dependent on repurposing a single, ancestral mechanosensory neuron type.

We report here our fortuitous discovery of cell type diversity of mechanosensory neurons in the tentacular ectoderm of the sea anemone *Nematostella vectensis,* which is a genome-enabled, genetically tractable cnidarian ^20–22^. *N. vectensis* gastrulates by invagination to form an embryo with ectoderm and endoderm separated by an extracellular matrix, the mesoglea ^23,24^. The embryo develops into a ciliated planula larva that swims with the aboral end oriented forward. The planula then undergoes life cycle transition, transforming into a polyp with oral tentacles whose ectoderm houses concentric hair cells with an apical cilium surrounded by a circle of 6-9 stereovilli ^14,25,26^. The apical sensory apparatus of the concentric hair cell is encircled by microvilli of adjacent support cells, forming a structure known as the hair bundle. As noted above, the multicellular structure consisting of a concentric hair cell surrounded by support cells is referred to as the hair bundle mechanoreceptor, and its mechanosensitivity has been confirmed by electrophysiology in *N. vectensis* ^14^. We have recently shown that the class IV POU transcription factor (POU-IV) – which is deeply shared across animals excepting Ctenophora ^27^ – regulates postmitotic differentiation and maturation of concentric hair cells by activating a specific set of effector genes (e.g. *polycystin-1* encoding a conserved transmembrane receptor of the TRP ion channel superfamily), and is necessary for touch-sensitivity of polyp tentacles ^26^. Upon further investigation into the anatomy and development of the tentacular nervous system in *N. vectensis,* we discovered a previously unrecognized population of concentric hair cells that are anatomically, molecularly and developmentally distinct from the conventional concentric hair cells of sea anemones, the evidence of which is detailed below.

## Results

### Sea anemone has two morphologically distinct types of hair cells

We first examined the morphological diversity of concentric hair cells in the ectodermal epithelium of oral tentacles in *N. vectensis* polyps. We used meganuclease-mediated transgenesis ^28^ to develop a *pou-iv* transgenic reporter line in which a *pou-iv* promoter (3.2 kb genomic sequence immediately upstream of the start codon of the *pou-iv* gene) drives the expression of photoconvertible fluorescent protein Kaede (Supplementary Fig. 1). Using this *pou-iv::kaede* transgenic line, we found that the *pou-iv* promoter drives Kaede expression in conventional concentric hair cells with a cilium (<15 μm in length; n=10) surrounded by a circle of 8-9 pronounced stereovilli (Figure 1A-E) - as previously reported using F0 *pou-iv::kaede* mosaic transgenic animals (Ozment et al., 2021). Unexpectedly, we also found Kaede expression in concentric hair cell-like cells with a longer erect cilium (<45 μm in length; n=13) and less pronounced stereovilli/microvilli (Figure 1F-J; Supplementary Movie 1). We will herein refer to the conventional concentric hair cell as the *type I hair cell*, and the morphologically distinct hair cell-like cell characterized by the long cilium as the *type II hair cell*.

**Figure 1:**
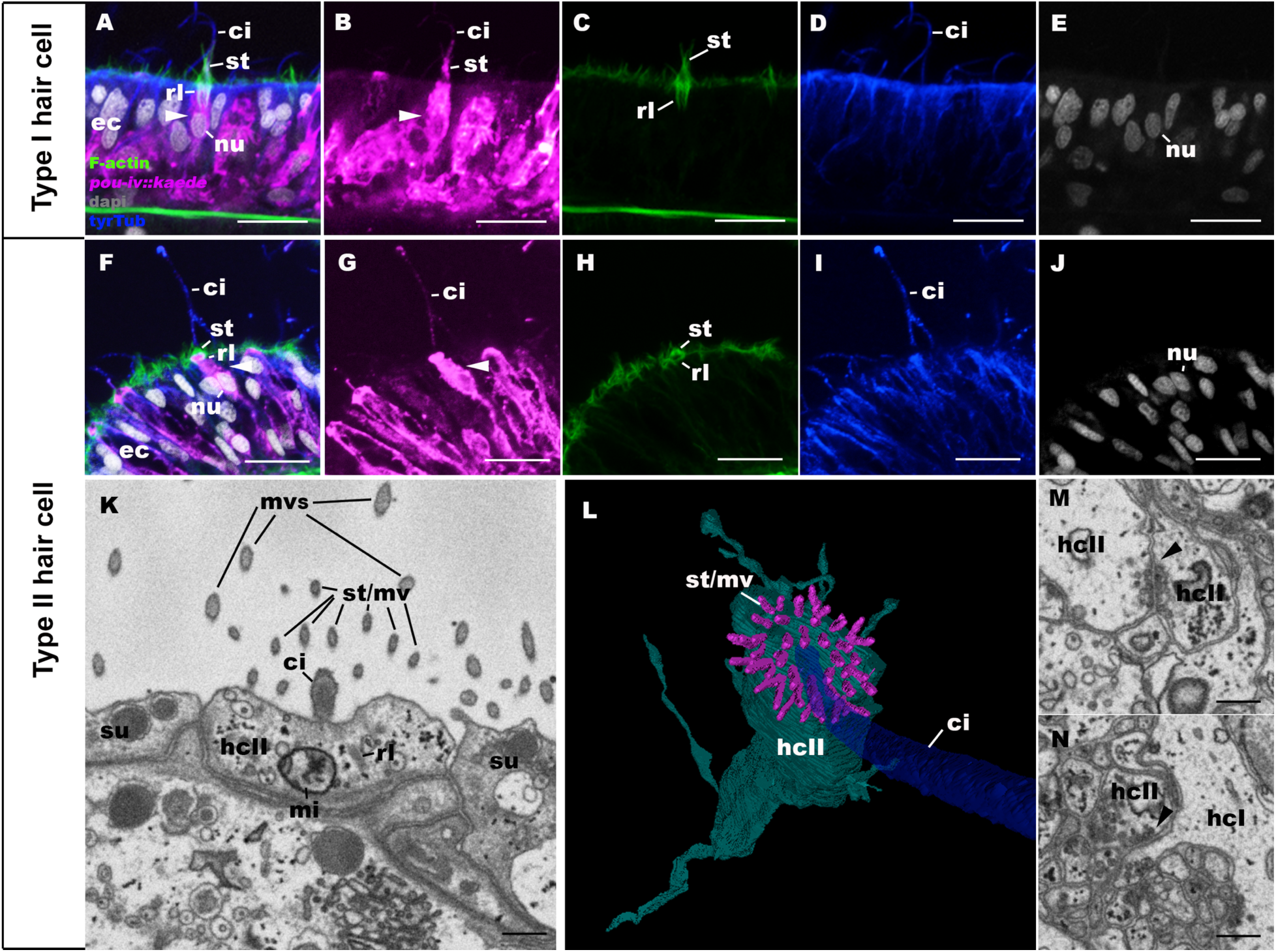
Tentacular ectoderm of the sea anemone *Nematostella vectensis* houses two morphologically distinct types of concentric hair cells. A-J: Confocal sections of the tentacular ectoderm (ec) of *pou-iv::kaede* transgenic *Nematostella vectensis* polyp, labeled with antibodies against Kaede (“pou-iv::kaede”) and tyrosinated ∂-tubulin (“tyrTub”). Filamentous actin is labeled with phalloidin (“F-actin”), and nuclei are labeled with DAPI (“dapi”). The apical epithelial surface faces up. Panels A-E show a type I hair cell (arrowhead), characterized by an apical cilium (ci) surrounded by stereovilli (st) with pronounced rootlets (rl). Panels F-J show a type II hair cell (arrowhead), characterized by a long cilium (ci) surrounded by short stereovilli (st) with rootlets (rl). K-N: serial block-face scanning electron microscopy (SBF-SEM) images and 3-D reconstruction of type II hair cells (hcII). Panel K shows the apical sensory apparatus consisting of a cilium (ci) surrounded by rings of microvilli (st/mv), a subset of which have actin rootlets that extend about 1 μm into the cytoplasm and therefore are stereovilli. Similar to type I hair cells (see ^14,26^), apical microvilli of adjacent support cells (mvs) encircle the apical mechanosensor of the type II hair cell. Panel L is an apical view of a 3-D reconstruction of a type II hair cell. Note that the cilium is surrounded by rings of about 30 stereovilli and microvilli (st/mv). Panels M and N show a two-way (bidirectional) chemical synapse between type II hair cells (arrowhead in M), and a one-way chemical synapse in which a type II hair cell is presynaptic to a type I hair cell (hcI) (arrowhead in N), respectively. These features are unique to type II hair cells; type I hair cells are not presynaptic to type I or type II hair cells. Note the clustering of dense-cored or opaque vesicles adjacent to parallel, relatively electron-dense plasma membranes. Abbreviations: nu nucleus; mi mitochondria Scale bar: 10 µm (A-J); 500 nm (K, M, N)

To better resolve the structure of the apical mechanosensory apparatus of the type II hair cell, we carried out serial block-face scanning electron microscopy (SBF-SEM). We discovered that the apical structure consisted of a single cilium surrounded by multiple rings of about 30 stereovilli and microvilli that were approximately 1 μm long (Figure 1K, L), contrasting stereovilli of the type I hair cell, which have been reported to range from 6-9 in number and 3-5 μm in length in *N. vectensis* ^14,26^.

Taking advantage of the SBF-SEM volume data, we next searched for chemical synapses of concentric hair cells in order to assess whether the pattern of synaptic connectivity would differ between type I and type II hair cells. Structurally, cnidarian chemical synapses consist of a pair of parallel plasma membranes with clear, dense-cored, or opaque vesicles - 70-150 nm in diameter - apposing one or both sides of the membranes, with cross filaments sometimes occurring in the intercellular space – the synaptic cleft – between the parallel plasma membranes (reviewed in ^29^). The SBF-SEM volume data enabled us to trace thin basal processes to the cell body and unambiguously determine the identity of the cell type to which each chemical synapse belonged. Indeed, we discovered conventional chemical synapses of both type I and type II hair cells in *N. vectensis*. Synaptic vesicles of type I and type II hair cells were observed to form distinct, dense clusters adjacent to rigid, parallel membranes sandwiching a synaptic cleft that is more electron-dense than the surrounding intercellular space (Figure 1M, N; Supplementary Fig. 2, 3; Supplementary Movie 2, 3). Synaptic vesicles of type I hair cells were dense-cored (Supplementary Fig. 3A-D), while those of type II hair cells were mostly opaque and sometimes dense-cored (Figure 1M, N; Supplementary Fig. 2, 3E-I; Supplementary Movie 2). Notably, synaptic vesicles of type I hair cells were significantly larger than those of type II hair cells (mean diameter: type I, 95.04 nm, n=17; type II, 70.38 nm, n=62; two-tailed t-test, p<0.001). In addition, the number of vesicles per synapse was found to be significantly lower in type I hair cells (mean: 13.5 per synapse; n=4 synapses) than in type II hair cells (mean: 88.3 per synapse, n=9 synapses) (two-tailed t-test, p<0.01). Chemical synapses of type I and type II hair cells were found primarily in neurites but sometimes also occur in the basal part of the cell body (i.e. basal to nucleus; see Supplementary Movie 4).

We then compared the pattern of synaptic connectivity in type I and type II hair cells. Both type I and type II hair cells were observed to form afferent/input synapses with epitheliomuscular cells (n=10 synapses found in type I hair cells; n=5 in type II), cnidocytes (nematocytes (n=2 in type I; n=1 in type II) and spirocytes (n=4 in type I; n=1 in type II)), and ganglion cells (n=3 in type I; n=8 in type II) (Supplementary Fig. 3 A-H). In addition, we found that type II hair cells, but not type I hair cells, formed two-way (bidirectional/reciprocal) synapses with each other (n=7; Figure 1M) and with ganglion cells (n=3; Supplementary Fig. 3I). Moreover, type II hair cells formed afferent synapses with type I hair cells (n=3; Figure 1N), but not vice versa in any of the type I hair cells that were examined (n=10 cells). These results indicate that type I and type II hair cells are distinct not only in the structure of apical mechanosensory apparatus, but also in the structure of synapses and the pattern of synaptic connectivity.

### Type I and type II hair cells are molecularly distinct

Next we asked whether the mechanotransduction mechanisms would differ between the two hair-cell types. As the first step towards tackling this question, we focused our attention on *polycystin-1* (PKD1) that encodes a transmembrane receptor of the Transient Receptor Potential (TRP) calcium channel superfamily. This gene was previously identified as a candidate mechanotransduction gene of the concentric hair cells, because it was found to be directly activated by POU-IV transcription factor specifically in hair cells ^26^. However, whether *polycystin-1* is expressed in type I hair cells, type II hair cells, or both, was unknown. We aimed to determine 1) whether *polycystin-1* was expressed in both types of hair cells, and 2) whether *polycystin-1* was essential for normal touch response behavior. To address whether *polycystin-1* is expressed in both types of hair cells, we have generated a stable transgenic reporter line for *polycystin-1* in which a *polycystin-1* promoter (2.1 kb genomic sequence immediately upstream of the start codon of the *polycystin-1* gene) drives the expression of Kaede photoconvertible fluorescent protein (Supplementary Fig. 4). We have confirmed by *in situ* hybridization that the expression pattern of *polycystin-1::kaede* (*pkd1::kaede*) indeed recapitulates that of endogenous *polycystin-1* at the primary polyp stage (Supplementary Fig. 5). Using this transgenic line, we found that *polycystin-1::kaede* reporter gene expression specifically occurred in type II hair cells, but not in type I hair cells, of tentacular ectoderm at the polyp stage (Figure 2A-D), evidencing that type I and type II hair cells are molecularly distinct.

**Figure 2:**
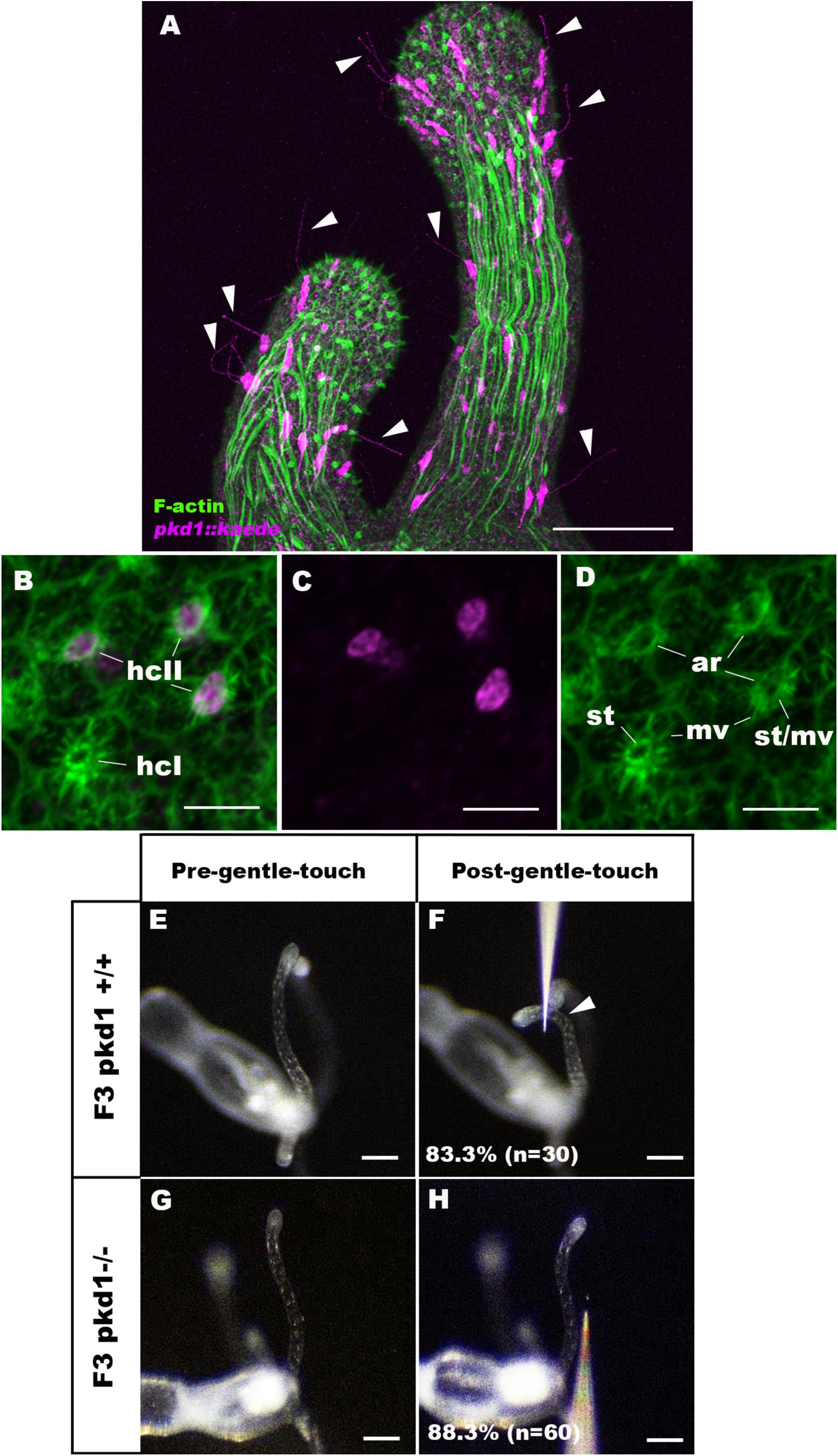
*polycystin-1* is a type-II-hair-cell-specific essential mediator of gentle touch response of oral tentacles in a sea anemone. A-D: Confocal sections of the tentacles of *pkd1::kaede* transgenic *Nematostella vectensis* polyp. Filamentous actin is labeled with SiR-Actin (A) or phalloidin (B, D) (“F-actin”), and Kaede is labeled with an anti-Kaede antibody in B and C. The distal end of the tentacle faces up in A. Panels B-D are sections at the level of the apical ectodermal epithelium. Arrowheads in A show long, often erect, cilia characteristic of type II hair cells. Panels B-D show *pkd1::kaede* expression in type II hair cells (hcII) with numerous thin stereovilli/microvilli (st/mv), but not in type I hair cells with discrete, large-diameter stereovilli (st). Note also the presence of a prominent contiguous F-actin ring (ar) lining the apicolateral membrane of *pkd1::kaede-*positive type II hair cells. Consistent with SBF-SEM data, adjacent support cells contribute microvilli (mv) to the apical mechanosensory apparatus of both types of hair cells. E-H: Behavior of wildtype (F3 *pkd1* +/+, E, F) and mutant (F3 *pkd1* −/−, G, H) *N. vectensis* polyps in response to gentle touch to their oral tentacles. Either a microinjection needle (shown) or a tungsten needle was brought close to the tentacle to elicit gentle touch response – a bending motion of the stimulated tentacle towards the source of the mechanical stimuli. Animals before (E, G) and after (F, H) gentle touch are shown. Most of wildtype animals displayed gentle touch response (83.3%, n=30; E, F), while the majority of *pkd1* homozygous mutants were touch-insensitive (88.3%, n=60; G, H). The arrowhead in F points to a stimulated tentacle exhibiting normal gentle touch response. Scale bar: 50 µm (A); 5 µm (B-D); 100 µm (E-H)

To assess whether type II-hair-cell-specific expression of *polycystin-1* is necessary for normal mechanosensory behavior in *N. vectensis,* we used CRISPR-Cas9-mediated mutagenesis to generate a *polycystin-1* mutant line in which the structure of TRP cation channel and the C-terminal cytoplasmic tail is disrupted (Supplementary Fig. 6). We predicted that if *polycystin-1* indeed encoded type II-hair-cell-specific mechanotransduction channels, perturbation of Polycystin-1 channel function should cause defects in type II hair cell function. Concentric hair cells with long, erect cilia – resembling type II hair cells of *N. vectensis* – occur in the ectoderm of tentacles in the hydroid *Coryne pintneri* ^4^, in the medusa of the hydrozoan *Aglantha digitale* ^9^, and in the polyp of the cubozoan *Carybdea marsupialis* ^12^. They are highly sensitive to gentle mechanical stimulation and water vibration; mechanical stimuli to the hair cell-bearing tentacle induce movement towards the source of the stimuli, involving bending of the stimulated tentacle in *Aglantha* and *Eutonina* hydrozoan medusae ^9,13^ and *Carybdea* polyp ^12^ and of the upper body of the polyp in *Coryne* ^4^. This behavior presumably facilitates the capture of nearby prey. In *Aglantha* medusae, this ‘pointing response’ is distinct from responses to stronger mechanical stimuli; direct contact of the tentacle triggers tentacular contraction and escape swimming behavior ^9^. In *N. vectensis* polyps, whole-tentacle contraction occurs in response to direct contact of a probe with the tentacular epithelium ^26^. We will here refer to this behavior as a *harsh touch response*. In addition, we have found that *N. vectensis* polyps display the pointing response; tentacles respond to an approaching probe, before the probe makes physical contact with the tentacular surface epithelium, by a quick bending motion that brings the tentacle closer to the object (see below). We will refer to this behavior as a *gentle touch response*. Given that type II hair cell-like cells with long, erect cilia in other cnidarians appear to be highly sensitive to gentle mechanical stimuli, we hypothesized that type II-hair-cell-specific expression of *polycystin-1* might be necessary for mediating the gentle touch response behavior in *N. vectensis*.

To test whether *polycystin-1* is essential for gentle touch response, we crossed *polycystin-1 +/−* heterozygous mutants (F1 or F2) with each other to generate a progeny with expected genotype frequencies of 25% −/− homozygotes, 50% +/− heterozygotes, and 25% +/+ wildtype siblings. We carried out gentle touch assay on the progeny, whereby a probe (a tungsten needle or a glass needle) is brought close (<1 mm) to a tentacle, without directly contacting the epithelial surface, to elicit bending of the tentacle towards the probe. We then separated the animals based on the presence/absence of the pointing response (Supplementary Movie 5), and genotyped them individually. We found significant enrichment of *polycystin-1* homozygous mutants in the gentle-touch-insensitive group (experiment 1, 55.3% (n=47), hypergeometric test p<0.001; experiment 2, 69.2% (n=39), p<0.001), and significant depletion of *polycystin-1* homozygous mutants in the gentle-touch-sensitive group (experiment 1, 10.6% (n=47), hypergeometric test p=0.012; experiment 2, 6.25% (n=32), p<0.01). Overall, only 11.7% (n=60) of *polycystin-1* homozygous mutants were deemed to exhibit gentle-touch response, while 83.3% (n=30) of their wildtype siblings showed normal gentle-touch response behavior (Figure 2E-H; Supplementary Movie 6, 7). In contrast, harsh-touch response occurred in *polycystin-1* homozygous mutants (100%, n=34; Supplementary Fig. 7, Supplementary Movie 8). We note that the gentle-touch-insensitive phenotype of *polycystin-1* homozygous mutants is not due to the failure to develop the apical sensory apparatus of type II hair cells (Supplementary Fig. 8). Neither did we find evidence that the phenotype was due to a change in the density of type II hair cells in *polycystin-1* homozygous mutants relative to wildtype animals; the estimated linear density of type II hair cells in *polycystin-1* homozygous mutants (mean: 14.2 per 100 µm unit length, n=5) did not significantly differ from that in their wildtype siblings (mean: 11.6 per 100 µm unit length, n=5) (two-tailed t-test, p = 0.5797). Taken together, these results strongly suggest that type II-hair cell-specific expression of *polycystin-1* is essential for gentle-touch behavior, but not for harsh-touch behavior, and that molecular mechanisms of mechanotransduction in type I and type II hair cells are not redundant.

### Type I and type II hair cells are developmentally distinct

We next considered whether type I and type II hair cells might represent different maturation phases of the same postmitotic cell type. For this possibility, two scenarios were conceivable; type I hair cells give rise to type II hair cells, or vice versa. The scenario whereby type II hair cells transform into type I hair cells seemed highly unlikely because *pkd1::kaede* expression specifically occurred in type II hair cells and not in type I hair cells (Figure 2A-D). If type II hair cells gave rise to type I hair cells, Kaede expression – stable enough to persist for months ^30^ – would be retained by type I hair cells, which was not observed in any F1 or F2 *pkd1::kaede* transgenic animals examined (n=15 animals). However, the possibility that type I hair cells transform into type II hair cells could not be ruled out.

To further explore the developmental relationship between type I and type II hair cells during maturation, we turned to the *pou-iv::kaede* transgenic line in which both type I and type II hair cells could be photoconverted and tracked. We photoconverted *pou-iv::kaede* primary polyps and examined whether the number of photoconverted type I hair cells would change relative to the number of photoconverted type II hair cells after 5 days of tentacular growth (Supplementary Fig. 9A, B). We predicted that if type I hair cells were the source of type II hair cells, the proportion of photoconverted type I hair cells relative to photoconverted type II hair cells should decrease as new type II hair cells formed. Conversely, if type II hair cells were the source of type I hair cells, the proportion of photoconverted type I hair cells relative to photoconverted type II hair cells should increase as new type I hair cells formed. The average proportion of *pou-iv::kaede-*positive type I relative to type II hair cells in tentacular ectoderm prior to photoconversion – assumed to correspond directly to the proportion immediately after photoconversion – was 1.04 (n=7 animals), and that of photoconverted type I hair cells relative to photoconverted type II hair cells after 5 days post-photoconversion was 1.00 (n=6 animals), failing to show any statistically significant change in either direction (two tailed t-test, p=0.876; Supplementary Fig. 9C). The number of *pou-iv::kaede-*positive type I or type II hair cells per unit volume did not significantly change either; the mean density of *pou-iv::kaede-*positive type I hair cells before photoconversion (101.26 per 10^6^ µm^3^; n=7 animals) did not significantly differ from that of *pou-iv::kaede-*positive photoconverted type I hair cells at 5 days after photoconversion (117.25 per 10^6^ µm^3^; n=6 animals) (two tailed t-test, p=0.501), and neither did the mean density of *pou-iv::kaede-*positive type II hair cells before photoconversion (98.24 per 10^6^ µm^3^; n=7 animals) significantly differ from that of *pou-iv::kaede-*positive photoconverted type II hair cells at 5 days after photoconversion (118.89 per 10^6^ µm^3^; n=6 animals) (two tailed t-test, p=0.069). The lack of evidence for transformation of type I hair cells into type II hair cells, or vice versa, is not due to insufficient time for completing the developmental process; we found type I and type II hair cells positive for non-photoconverted Kaede (Supplementary Fig. 9D-I), which indicates that the incubation period was sufficient for generating both types of hair cells anew. We therefore do not find robust support for the hypothesis that type I and type II hair cells represent different maturation states of the same postmitotic cell type.

If type I and type II hair cells indeed develop from distinct postmitotic precursors, the pattern of development may also differ between the two cell types. To investigate this possibility, we compared the timing of when type I and type II hair cells would begin postmitotic differentiation, taking advantage of the *pou-iv::kaede* transgenic reporter line in which postmitotic precursors to hair cells, as well as other cell types (e.g. cnidocytes), could be labeled by photoconversion and tracked (Supplementary Fig. 10, 11). Morphologically and/or molecularly differentiated forms of type I and type II hair cells become evident at the tentacle-bud stage (type I, ^26^; type II, Supplementary Fig. 4), and therefore, early postmitotic precursors to type I and type II hair cells are expected to form at or prior to the tentacle-bud stage. To test whether postmitotic precursors to type I and/or type II hair cells emerge during planula development, we photoconverted *pou-iv::kaede*-positive cells at different stages of planula development (2 dpf early planula, 3 dpf mid-planula I, 4 dpf mid-planula II, and 5 dpf late planula; staging based on ^25^), and followed their fate at the tentacle-bud and primary polyp stages (9-12 dpf) (Figure 3). When photoconversion was carried out at the early planula stage, we did not find photoconverted type I or type II hair cells in the tentacular ectoderm of primary polyps, but observed photoconverted cnidocytes (Figure 3F-H; early planula, n=5 animals). This confirms that postmitotic precursors to hair cells develop post-embryonically, and suggests that at least a subset of embryonically generated cnidocytes become incorporated into tentacles of primary polyps. When photoconversion was performed at the mid-planula I stage, a subset of morphologically unambiguous type I hair cells, but not type II hair cells, were occasionally found to be photoconverted (Figure 3I-K; n=5 animals). Photoconverted type I hair cells were consistently found in animals in which photoconversion occurred at the mid-planula II stage (n=7 animals). This suggests that postmitotic precursors to type I hair cells emerge between early planula and mid-planula II stages (i.e. between 48-96 hpf at room temperature). Photoconverted type II hair cells with mature morphology, on the other hand, were consistently observed at the tentacle-bud/primary polyp stage in animals where photoconversion was performed at the late planula stage (Figure 3L-N; n=7 animals), indicating that postmitotic precursors to type II hair cells begin to develop between mid-planula II and late planula stages (i.e. between 96-120 hpf at room temperature). Taken together, these results suggest that early postmitotic precursors to type I hair cells form before those to type II hair cells during planula development, indicative of temporally distinct generative mechanisms for type I and type II hair cells.

**Figure 3:**
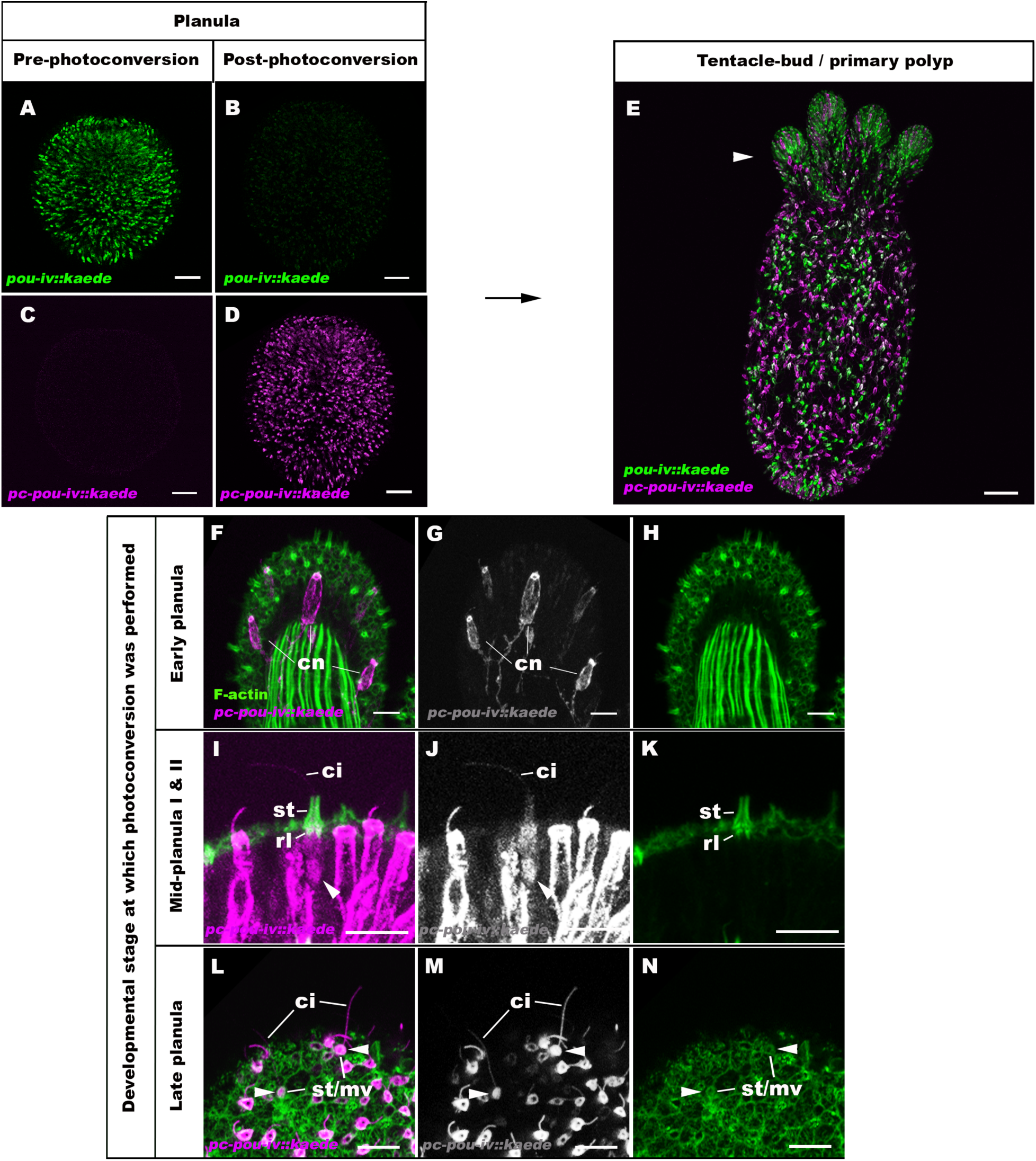
Type I and type II hair cells have temporally distinct developmental origins in the sea anemone *Nematostella vectensis*. Confocal sections of *pou-iv::kaede* transgenic *N. vectensis* at the planula (A-D) and tentacle-bud/primary polyp (E-N) stages. Kaede fluorescent protein (“*pou-iv::kaede*”) was photoconverted (“*pc-pou-iv::kaede*”) by ultraviolet illumination during planula larval development (e.g. A-D), and photoconverted Kaede was used as a cell marker to trace the fate of *pou-iv::kaede*-positive larval cells at life cycle transition (E-N). Filamentous actin (“F-actin”) is labeled with a fluorescent dye SiR-Actin. A-E show sections through the entire animal with the blastopore/mouth facing up. F-H are sections through the center of a tentacle with photoconverted cnidocytes (cn); the oral side of the tentacle faces up. Photoconversion was performed at the early planula stage (2 dpf). Photoconverted type I and type II hair cells were absent, confirming their post-embryonic origin. I-K show sections through a photoconverted type I hair cell having characteristically pronounced stereovilli (st) with actin rootlets (rl) in an animal where photoconversion was performed at the mid-planula I stage (3 dpf). The apical epithelial surface faces up. Photoconverted type II hair cells were absent, implying that early-born type I and type II hair cells have temporally distinct developmental origins. L-N show sections at the level of the surface epithelium of the tentacle primordium of an animal in which photoconversion was performed at the 5 dpf late planula stage. Note the presence of photoconverted type II hair cells with characteristically long cilia (>20 µm; ci), indicating that type II hair cell precursors begin to develop after type I hair cell precursors. Abbreviations: st/mv rings of stereovilli/microvilli. Scale bar: 50 µm (A-F, J, N); 10 µm (G-I, K-M, O-Q)

Postmitotic differentiation and maturation of both type I and type II hair cells are dependent on POU-IV, evidenced by the loss of long stereovillar rootlets and *polycystin-1* expression – defining characteristics of type I and type II hair cells, respectively – in *pou-iv* null mutant polyps ^26^. However, the role of POU-IV in specifying and/or maintaining the identity of these cell types may differ. To explore the possibility that *pou-iv* expression is differentially regulated in the two cell types, we used the *pou-iv::kaede* transgenic reporter line to compare the degree of persistence of *pou-iv* promoter activity during the maturation of type I and type II hair cells. We photoconverted *pou-iv::kaede-*positive, immature type I and type II hair cells at the late planula stage, and allowed the photoconverted animals to develop into primary polyps. We then examined whether photoconverted type I and type II hair cells expressed non-photoconverted Kaede, in order to determine whether the *pou-iv* promoter activity would persist during maturation. We found that the majority of the photoconverted type I hair cells did not express non-photoconverted Kaede at detectable levels (66.7%, n=27 cells across 6 animals; Supplementary Fig. 12A-C), suggesting that *pou-iv* promoter activity in type I hair cells is transient and does not persist through the maturation process. In contrast, photoconverted type II hair cells always expressed non-photoconverted Kaede at levels comparable to those of newly-born type II hair cells (n=14 cells across 6 animals; Supplementary Fig. 12D-F), implying that *pou-iv* promoter activity persisted in type II hair cells throughout the maturation process.

Next, we asked whether *pou-iv* promoter activity persisted in mature type II hair cells. To address this question, we photoconverted *pou-iv::kaede-*positive cells at the primary polyp stage, to label morphologically mature type II hair cells along with other *pou-iv::kaede-*positive cells, and allowed the photoconverted animals to continue growth for 5 days. We found that all of the photoconverted type II hair cells that were examined expressed non-photoconverted Kaede (n=51 cells across 7 animals; Supplementary Fig. 12G-I). Assuming that at least a subset of photoconverted type II hair cells were at a mature state at the time of photoconversion, the results suggest that *pou-iv* promoter activity continues in mature type II hair cells. Taken together, these findings indicate that *pou-iv* promoter activity is differentially regulated in type I and type II hair cells, and raise the possibility that the transcriptional regulatory mechanism that establishes and maintains mature cell identity differs between type I and type II hair cells.

## Discussion

Here we have presented anatomical, molecular and developmental evidence for cell type diversity in cnidarian mechanosensory neurons. Specifically, we uncovered in the tentacular ectoderm of the sea anemone *N. vectensis* previously unrecognized mechanosensory neurons – type II hair cells – that differ in morphology, mechanotransduction mechanism, and development from the conventional mechanosensory neurons of the sea anemone – the type I hair cells. Notably, we have not only identified, for the first time, classical chemical synapses in cnidarian hair cells, but also discovered that type I and type II hair cells have different synaptic structures and patterns of synaptic connectivity. In addition, we have reported the first ion channel-encoding gene – *polycystin-1 -* essential for mechanosensory behavior in Cnidaria, or any non-Bilateria, and have found that this gene mediates gentle touch response specifically in type II hair cells and not in type I hair cells, consistent with distinct mechanotransduction mechanisms being deployed in the two cell types. Also noteworthy is the first experimental determination of temporal developmental origins of postembryonic cell types in Cnidaria, and the discovery that postmitotic precursors to type I and type II hair cells begin to emerge at different phases of larval development, indicative of temporally distinct developmental mechanisms.

The molecular mechanisms of development and function of the newly discovered type II hair cells in the sea anemone appear similar to those of mechanosensory cells in Bilateria, indicative of deeply shared ancestry. For instance, the class IV POU homeodomain transcription factor regulates differentiation not only of type II hair cells in the sea anemone *N. vectensis* ^26^, but also of mechanosensory neurons/cells across bilaterians (e.g. worms and vertebrates ^31–36^). Moreover, in *N. vectensis,* effector genes that are directly activated by POU-IV in type II hair cells – corresponding to a transcriptomically defined adult cell type uniquely expressing *polycystin-1* (‘metacell c79’; ^37^) – show significant enrichment of the GO term ‘sensory perception of sound’ ^26^. Given that evidence for this GO annotation derives from bilaterian data, it suggests that the effector genes that define the mechanoreceptor identity are broadly shared across the cnidarian type II hair cell and bilaterian mechanosensory cells. Of note, in the hydrozoan cnidarian *Hydra vulgaris, pou-iv* and *polycystin-1* are co-expressed in a specific cluster of transcriptomically similar neurons (‘ec1’; ^38^), concordant with the core gene regulatory network for type II hair cell development being conserved across Cnidaria. It will be important to examine whether *pou-iv* and *polycystin-1* are indeed involved in the development of mechanosensory neurons in *Hydra*; specialized mechanosensory neurons have not been identified in this cnidarian. Intriguingly, the gene regulatory mechanism for mechanoreceptor development involving *pou-iv* and *polycystin-1* might even predate the divergence of the Cnidaria/Bilateria clade and its likely sister group Placozoa (^39–42^; but see ^43,44^ for an alternative view of phylogeny), as unambiguous orthologs of *pou-iv* and *polycystin-1* are not only present in the Placozoa ^26,27^ but are co-expressed in a transcriptomically defined peptidergic cell type (‘metacell 214’ in *Trichoplax sp. H2*; ^45^). Investigation of whether these *polycystin-1-*positive peptidergic cells represent specialized mechanosensory cells of Placozoa is therefore warranted. Taken together, current comparative evidence indicates that the molecular mechanisms of development and function of type II hair cells are not sea anemone lineage-specific, but have deeper evolutionary roots.

Our findings suggest that anatomy, development, and mechanotransduction mechanisms of cnidarian mechanosensory neurons are diverse. As described above, within-individual morphological variation in apical mechanosensors of presumptive mechanosensory neurons occurs across Cnidaria (e.g. ^6,9,10,12,17,46^), and therefore, it is plausible that diverse types of mechanosensory neurons exist beyond sea anemone cnidarians. In addition, the planula larvae – the post-gastrulation dispersal phase conserved across Cnidaria – are thought to be mechanosensitive (e.g. geotactic, ^47,48^; phonotactic, ^49–51^), although the cellular bases of mechanosensation remain unknown. If mechanosensory neurons indeed existed in *N. vectensis* planulae, they would be distinct from type I and type II hair cells, as mature forms of type I and type II hair cells are absent in planulae. We suggest that cell type diversity of cnidarian mechanosensory neurons across life cycle stages and lineages likely remains underestimated and merits further investigation.

Our discovery of structurally, molecularly and developmentally distinct mechanosensory neurons from a cnidarian further implies that evolutionary histories of mechanosensory neurons in non-bilaterian animals are not simple. Repurposing of a single type of mechanosensory neurons as the sole mechanism of mechanosensory neural evolution is no longer tenable for Cnidaria. Instead, cell type diversification of mechanosensory neurons must have occurred in Cnidaria – paralleling mechanoreceptor evolution in Bilateria – and/or in early animal ancestors basal to Cnidaria and Bilateria followed by the retention of ancestrally distinct mechanosensory neuron types in the descendant lineages. Resolving the resultant alternative hypotheses – independent cell type diversification in Cnidaria and Bilateria, ancient radiation predating the divergence of Cnidaria and Bilateria, or a combination of both – necessitates a deeper understanding of the mechanisms of development and function of distinct mechanosensory neuron types within Cnidaria and across deeply-branching animal lineages.

## Methods

### Animal culture

*Nematostella vectensis* were cultured as previously described ^52,53^.

### Generation of *kaede* transgenic lines

The *pou-iv::kaede* and *pkd1::kaede* transgenic animals were produced by I-SceI-mediated transgenesis as described previously ^26,28^ with modifications. To generate *pou-iv::kaede* plasmid, 3199 bp genomic sequence upstream of the start codon of the *Nematostella vectensis pou-iv* (*Nematostella vectensis v1.0*; scaffold 16: 1065408-1068606; https://mycocosm.jgi.doe.gov/Nemve1/Nemve1.home.html) was cloned in front of the open reading frame of the Kaede gene ^30^ by FastCloning ^54^. To generate *pkd1::kaede* plasmid, 2145 bp genomic sequence upstream of the start codon of the *Nematostella vectensis polycystin 1* (scaffold 353: 49524-51668; https://mycocosm.jgi.doe.gov/Nemve1/Nemve1.home.html), was cloned in front of the open reading frame of the Kaede gene. The plasmid was digested with I-SceI for 15-30 minutes at 37 °C and injected into zygotes at 50 ng/ μl. The injected animals that were identified as Kaede-positive (F0 animals) were raised to sexually mature adult polyps. F0 animals were then crossed with each other to generate F1 progeny, which was screened to identify carriers. These Kaede-positive F1 animals were raised to adult polyps, which were individually crossed with wildtype animals. In all cases examined, approximately half of the F2 progeny showed Kaede fluorescence, consistent with F1 animals being heterozygous. Both F1 and F2 animals were used in this study.

### CRISPR-Cas9-mediated mutagenesis

20 nt-long sgRNA target sites were manually identified in the *N. vectensis polycystin-1* genomic locus (*Nematostella vectensis v1.0*; scaffold 353:51565-82045; https://mycocosm.jgi.doe.gov/Nemve1/Nemve1.home.html), specifically in the region that encodes the TRP ion channel (scaffold 353:76524-80526). To minimize off-target effects, target sites that had 17 bp-or-higher sequence identity elsewhere in the genome were excluded.

Selected target sequences were as follows.

5’- CCCAGTCGTAGAAATTCTCG-3’ (Cr1)

5’- TTGTCCATAACTGTAAGACT-3’ (Cr2)

5’- TTCCTCTGTCTGACCCAGCT-3’ (Cr3)

5’- ATGTTGACCAGGACCCTGAA-3’ (Cr4)

The sgRNA species were synthesized *in vitro* (Synthego) and mixed at equal concentrations. The sgRNA mix and Cas9 endonuclease (PNA Bio, PC15111, Thousand Oaks, CA, USA) were co-injected into fertilized eggs at concentrations of 500 ng/µl and 1000 ng/µl, respectively.

### Genotyping of embryos and polyps

Genomic DNA from single embryos or from tentacles of single polyps was extracted by using a published protocol ^22,55,56^, and the targeted genomic locus was amplified by nested PCR. Primer sequences used for nested genomic PCR are: “1” Forward 5’-GAGTGCGTTCTTTCGATTCGGTGAG-3’, “1” Reverse 5’- GGCAAATACGTCCATGATAATCGTC-3’, “2” Forward 5’- TGCTATCATTGATGCTGTTCCAGTGC-3’, “2” Reverse 5’- AATCGGACCCAAGATCGGCTGCG-3’. To determine the sequence of mutant alleles, PCR products from genomic DNA extracted from F1 mutant polyps were gel-purified, cloned and sequenced using a standard procedure. Using the sequence information of the *polycystin-1-* mutant allele, genotyping primers for F2/F3 animals were designed as follows (Supplementary Fig. 6B).

Forward 5’- GACAATTTCTGTAATGTGACGTGACC-3’

Reverse (1), 5’- CAGTGAAGCCCACGTCGTACG-3’ (*polycystin-1+* -specific; expected size of PCR product, 249 bp)

Reverse (2), 5’- GAAGAAGACAAGGAAGACCACAGAG-3’ (expected size of PCR product: *polycystin-1+*, 3554 bp; *polycystin-1-*, 829 bp)

### Behavioral analysis

10dpf or older, unfed primary polyps with extended tentacles were used for behavioral analyses. For gentle touch assay, a tungsten needle attached to a microdissection needle holder (Roboz Surgical Instrument Co., Gaithersburg, MD, USA), or a microinjection glass needle attached to a micromanipulator (MO-202U; Narishige), was brought close (<1 mm) to a tentacle without direct contact with the epithelium. The response of 2-4 tentacles per individual polyp was examined. The polyp that bent its tentacle(s) towards the needle was deemed gentle-touch-sensitive. For harsh touch assay, the needle was moved to directly contact the surface epithelium of the tentacle. The polyp that contracted the stimulated tentacle was deemed harsh-touch responsive. Behavioral experiments were performed and recorded under a Zeiss Stemi 508 microscope or a SteREO Discovery V8 microscope equipped with Nikon DSL-4 camera. Recorded movies were viewed and annotated by using Image J.

### Live imaging

Specimens were transferred from 1/3 seawater solution to a solution of 2.43% MgCl_2_ in 1/3 seawater to anesthetize for 15 minutes at room temperature. Specimens and MgCl_2_ solution were placed in chamber slides (Lab-Tek II Chambered Coverglass W/Cover #1.5 Borosilicate Sterile, Nalge Nunc International #155360), and imaged using a Zeiss LSM 900 or Nikon Ti2 inverted microscope equipped with Nikon DSL-4 camera. Images were viewed using Image J or NIS Elements Ar (ver 5.11.01).

To estimate the linear density of type II hair cells, the number of characteristically long erect cilia (> 20 µm) – assumed to belong to type II hair cells – projecting perpendicular to the surface ectoderm was measured per unit length (100 µm) of the distal portion of a tentacle (i.e. from the distal epithelial tentacle tip extending 100 µm proximally toward the body column). Nikon NIS Elements Advanced Research software (ver. 5.11.01) was used to capture and annotate images for cell counts.

### Kaede photoconversion

Kaede fluorescence was converted from green to red by a 385 nm wavelength violet light using a X-Cite XYLIS LED illumination System (XT720L; Excelitas). Prior to photoconversion, *pou-iv::kaede* transgenic animals were mounted on a microscope slide in order to immobilize them. Transgenic animals were then exposed to a 385 nm wavelength LED light for 1 minute at 100% power using a 10x objective on a Zeiss LSM900.

### Immunofluorescence

Animal fixation and immunohistochemistry were performed as previously described ^25,57^. For immunohistochemistry, we used primary antibodies against POU-IV (rabbit, 1:200; Ozment et al., 2021), Kaede (rabbit; 1:500; Medical & Biological Laboratories, PM012M), and tyrosinated ∂-tubulin (mouse, 1:500, Sigma T9028), and secondary antibodies conjugated to AlexaFluor 568 (1:200, Molecular Probes A-11031 (anti-mouse) or A-11036 (anti-rabbit)) or AlexaFluor 647 (1:200, Molecular Probes A-21236 (anti-mouse) or A-21245 (anti-rabbit)). Nuclei were labeled using the fluorescent dye DAPI (1:1,000, Molecular Probes D1306), and filamentous actin was labeled using AlexaFluor 488-conjugated phalloidin (1:25, Molecular Probes A12379) or SiR-Actin (1:1000, Cytoskeleton, Inc. CY-SC001). We note that anti-Kaede immunoreactivity occurs in a subset of endodermal neurons (Supplementary Fig. 13), indicating that these neurons express Kaede-like proteins. This cross-reactivity makes it difficult to ascertain Kaede expression in endoderm when immunostaining is performed with an anti-Kaede antibody; we therefore avoided the use of the anti-Kaede antibody for analyses of Kaede expression in the endoderm. Specimens were mounted in ProlongGold antifade reagent (Molecular Probes, P36930) or Vectashield antifade reagent (Vector Laboratories H-1000-10). Fluorescent images were recorded using a Zeiss LSM900. Images were viewed using ImageJ.

### Double fluorescent situ hybridization with immunofluorescence

Animal fixation was performed as previously described ^25,57^. The double fluorescent in situ hybridization procedure was modified from a previously described protocol ^25^. Fixed *polycystin-1::kaede* transgenic animals were washed in PBST solution (0.1% Tween20 in 1X PBS) then underwent digestion with Proteinase K (final concentration = 0.02 mg/mL), followed by glycine (2 mg/mL) washes, triethanolamine (1%, pH 7.8) washes, and addition of acetic anhydride. Next, samples were washed again in PBST and were refixed in a 4% solution of paraformaldehyde, followed by PBST washes. Specimens were incubated with hybridization buffer for the prehybridization step. Probes were added at a final concentration of 1 ng/µL and left to incubate for 65 hours at 60°C. An antisense digoxigenin-labeled riboprobe against *N. vectensis polycystin 1* was synthesized from 3’ RACE products, and antisense fluorescein-labeled probes were generated against *kaede* (MEGAscript transcription kit; Ambion). After 65 hours, samples were incubated in fresh hybridization buffer and then underwent posthybridization washes consisting of increasing concentrations of 2X SSC (pH 7), then 0.05X SSC, and finally PBST again. Specimens were blocked in blocking buffer (0.5% blocking reagent (Perkin Elmer) in PBST) and then underwent overnight incubation with 1:100 Anti-Digoxigenin-POD (Roche, REF 11633716001) and 1:1000 acetylated ∂-tubulin (mouse, 1:500, Sigma T6793). Samples were washed again in PBST, then incubated with fluorophore tyramide amplification reagent (HCA ImagAmp 546 Kit NEL774001KT, Revvity; or HCA ImagAmp 647 Kit NEL775001KT, Revvity). After incubation, specimens were washed in 3% H_2_O_2_ in PBST, followed by a PBST only wash and blocking in blocking buffer. Specimens were incubated overnight in 1:100 Anti-Fluorescein-POD (Roche, REF 11426346910) and 4:1000 AlexaFluor 568 anti-mouse (Invitrogen A-11031) or AlexaFluor 647 anti-mouse (Invitrogen A-21236). After incubation samples were washed in PBST and then incubated with fluorophore tyramide amplification reagent (HCA ImagAmp 488 Kit NEL771001KT, Revvity). Samples were washed again in PBST then nuclei were labeled using the fluorescent dye DAPI (1:1,000, Molecular Probes D1306. Specimens were mounted in ProlongGold antifade reagent (Molecular Probes, P36930). Fluorescent images were recorded using a Zeiss LSM900. Images were viewed using ImageJ.

### EdU pulse labeling

*Pou-iv::kaede* transgenic animals were incubated in 1/3 seawater containing 10 μM of the thymidine analogue, EdU (Click-iT EdU AlexaFluor 488 cell proliferation kit, C10337, Molecular Probes), for 20 minutes to label S-phase nuclei. Following washes in fresh 1/3 seawater, the animals were immediately fixed as described previously ^25,57^. Immunohistochemistry was then carried out as described above. Following the immunohistochemistry procedure, fluorescent labelling of incorporated EdU was conducted according to the manufacturer’s recommendations prior to DAPI labelling.

### Serial block-face scanning electron microscopy (SBF-SEM) and data handling

Sample preparation for SBF-SEM was carried out based on a previously established protocol ^58^ with modifications. 10 dpf primary polyps were anesthetized in 2.43% MgCl_2_ for 20 minutes, and then fixed in 2.5% glutaraldehyde in 0.1 M sodium cacodylate buffer (pH 7.4) containing 2mM CaCl_2_ at 4°C overnight. Fixed polyps were washed in 0.1 M sodium cacodylate buffer containing 2mM CaCl_2_ for 10 min on ice. They were then washed in 0.1 M sodium cacodylate buffer containing 2mM CaCl_2_ and 50 mM glycine for 10 min on ice. Samples were subsequently rinsed in 0.1 M sodium cacodylate buffer containing 2mM CaCl_2_ for 10 minutes on ice twice, and were kept overnight at 4°C. Samples were incubated in 2% osmium tetroxide buffered in 0.1 M sodium cacodylate for 45 minutes at room temperature (RT). The osmium solution was replaced with 1.5% potassium ferrocyanide in 0.1 mM cacodylate buffer, in which samples were incubated for 45 minutes at RT in the dark. Samples were washed with water for 10 minutes twice. Samples were incubated in 1% thiocarbohydrazide in water for 15 minutes at 40 °C, and were rinsed in water for 10 minutes twice. They were then incubated in 1% osmium tetroxide for 45 minutes at RT, and washed in water for 10 minutes twice. Samples were incubated in 1% uranyl acetate in 25% ethanol for 20 minutes at RT in the dark, and were washed in water for 5 minutes three times and were left overnight at 4°C. Next, samples were stained with Walton’s lead aspartate for 45 minutes at 60°C, and were dehydrated in a graded ethanol series (30%, 50%, 70%, 90%, 100%), 1:1 ethanol to acetone, and then 100% acetone. Samples were then infiltrated with 3:1 acetone to Durcupan resin (Sigma-Aldrich) for 2 hours, 1:1 acetone to resin for 2 hours, 1:3 acetone to resin overnight, and then flat embedded in 100% resin on a glass slide and covered with a sheet of Aclar at 60°C for 48 hours. The resin was shaved to expose the polyp tentacle surface using an ultramicrotome (UC7, Leica), and was remounted to a metal pin with conductive silver epoxy (CircuitWorks, Chemtronics). A polyp tentacle was sectioned and imaged at the Electron Microscopy Core Facility at Max Planck Florida Institute for Neuroscience using 3View2XP and Digital Micrograph (ver. 3.30.1909.0, Gatan Microscopy Suite) on a Gemini SEM300 (Carl Zeiss Microscopy LLC.) equipped with an OnPoint BSE detector (Gatan, Inc.) and Focal Charge Compensation module (Carl Zeiss Microscopy LLC.). Imaging was performed at 2 kV accelerating voltage, 20 um aperture, 1.2 us pixel dwell time, 30% nitrogen gas flow, 5.149 nm per pixel, and 35.173 nm section thickness. The detector magnification was calibrated within SmartSEM imaging software (ver. 6.0, Carl Zeiss Microscopy LLC.) and Digital Micrograph with a 500 nm cross-line grating-standard following sample acquisition using the same imaging parameters. True Z-section thickness was calculated using the cylindrical diameters method ^59^ on longitudinal sections through microvilli. Serial images were exported as TIFFs to TrakEM2 ^60,61^ in ImageJ (ver. 1.52p) and aligned using the Scale-Invariant Feature Transform algorithm with linear feature correspondences and rigid transformation (Lowe, 2004). A type II hair cell and synapses were reconstructed in 3D by using segmentation tools and the software ImageJ 3D Viewer ^62^ in TrakEM2. Image analysis tools in TrakEM2 were used to measure the diameter of synaptic vesicles and the total number of synaptic vesicles per synapse. To estimate the total number of synaptic vesicles per synapse, vesicle clouds – defined as a cluster of at least 3 vesicles within about 100 nm of each other (cf. ^63^) – located at each synapse were segmented. The area of a vesicle cloud was summed across a series of sections for each synapse and was multiplied by the section thickness (35.173 nm) to obtain the total volume of the vesicle cloud. In addition, for each synapse, we measured the diameter of individual vesicles (n=2 or more per synapse) and calculated the mean diameter, which was used to estimate the expected volume of a vesicle for the synapse. The total volume of the vesicle cloud was then divided by the expected volume of a synaptic vesicle to estimate the total number of synaptic vesicles for each synapse.

## Data Availability

Source Data are provided with this paper.

## Acknowledgements

We thank Ethan Ozment for fixing samples for electron microscopy, and Kailey Hall for animal care. We are also grateful to Connon Thomas and Naomi Kamasawa at the Imaging Center and Electron Microscopy Core Facility at Max Planck Florida Institute for Neuroscience for their assistance with generating Serial Block-Face Scanning Electron Microscopy data. Lastly, we would like to thank the three anonymous reviewers for their constructive feedback on an earlier version of the manuscript, which helped improve the manuscript. This work was supported by Arkansas Bioscience Institute, University of Arkansas, and National Science Foundation Grant No.1931154 and 2042529.

## Author Contributions

Conceptualization: N.N. Investigation: J.B., K.M., M.A.P.S., S.R., G.S., N.N. Data analysis: J.B., N.N. Methodology: J.B., N.N. Funding acquisition: N.N. Supervision: N.N. Visualization: J.B., N.N. Writing: J.B., N.N.

## Competing interests

No competing interests declared.

## SUPPLEMENTARY INFORMATION

**Supplementary Movie 1: A long, erect cilium of a type II hair cell**

A 3D projection image of confocal sections through the surface ectoderm of a live tentacle of pou-iv::kaede transgenic adult polyp, rotated around the y-axis. pou-iv::kaede expression is shown in grey. The purple arrowhead points to a <50 µm long, erect cilium characteristic of type II hair cells. Scale bar: 50 µm

**Supplementary Movie 2: A serial z-stack SBF-SEM images of a two-way chemical synapse between type II hair cells shown in Figure 1M and Supplementary Fig. 2A**

Scale bar: 500 nm

**Supplementary Movie 3: A 3-D model of a two-way chemical synapse between type II hair cells shown in Supplementary Fig. 2B, rotated around the y axis**

**Supplementary Movie 4: A lateral view of a 3-D reconstruction of a type II hair cell shown in Figure 1L, rotated around the y-axis.**

The apical side is up. The cilium – the apical projection – is shown in blue, stereovilli/microvilli in purple, the cell body and basal neurites in cyan, the nucleus within the cell body in blue, and vesicle clouds associated with synapses in yellow. Note that vesicle clouds, and therefore synapses, are located primarily in basal regions including neurites.

**Supplementary Movie 5: Gentle touch assay**

A movie of gentle touch assay where a tungsten needle was brought close to F3 *polycystin-1* primary polyps with the expected frequencies of 25% *polycystin-1* +/+, 50% *polycystin-1* +/−, and 25% *polycystin-1* −/− genotypes. Polyps that bent their tentacles towards the needle were deemed gentle-touch-sensitive (green arrowhead), while those that did not were deemed gentle-touch-insensitive (purple arrowhead).

**Supplementary Movie 6: Gentle touch assay on an F3 *pkd1* +/+ *N. vectensis* polyp**

**Supplementary Movie 7: Gentle touch assay on an F3 *pkd1* −/− *N. vectensis* polyp**

**Supplementary Movie 8: Harsh touch assay on an F3 *pkd1* −/− *N. vectensis* polyp**

**Supplementary Figure 1:**
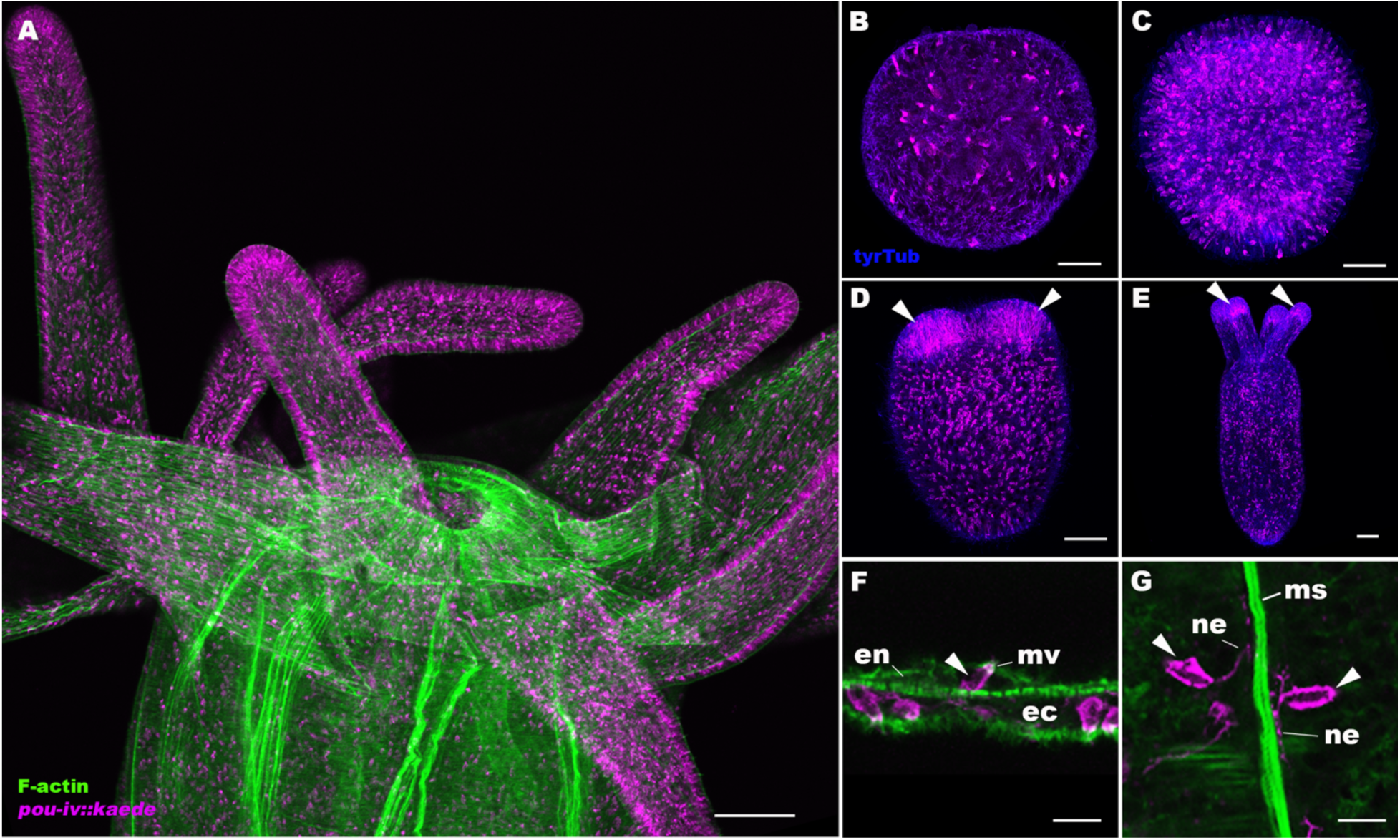
Developmental expression pattern of Kaede fluorescent protein in *pou-iv::kaede* transgenic *Nematostella vectensis*. Confocal sections of *pou-iv::kaede* transgenic *N. vectensis* at juvenile polyp (A), gastrula (B), planula (C), tentacle-bud (D), and primary polyp (E-G) stages, labeled with an antibody against tyrosinated ∂-tubulin (“tyrTub”). Filamentous actin is labeled with phalloidin (A) or SirActin (F, G) (“F-actin”), and Kaede is labeled with an anti-Kaede antibody in A-E. Panels A-E show side views of animals with the blastopore/mouth facing up. Panels A and E show sections through the entire animal, and panels B-D show sections through surface ectoderm. Panel F shows sections through a cnidocyte (arrowhead) in the endoderm of the body column, with the apical microvilli (mv) facing up. Panel G shows sections through endodermal cnidocytes (arrowheads) extending basal neurite-like processes (ne) along the longitudinal muscle fibers (ms); the oral side is up. *pou-iv::kaede* is expressed predominantly in the ectoderm in scattered epithelial cells throughout development; in the endoderm, *pou-iv::kaede* is expressed in cnidocytes, which are rare. Arrowheads in D and E indicate concentration of *pou-iv::kaede* expression in the tip of growing tentacles. Scale bar: 100 µm (A); 50 µm (B-E); 10 µm (F, G)

**Supplementary Figure 2:**
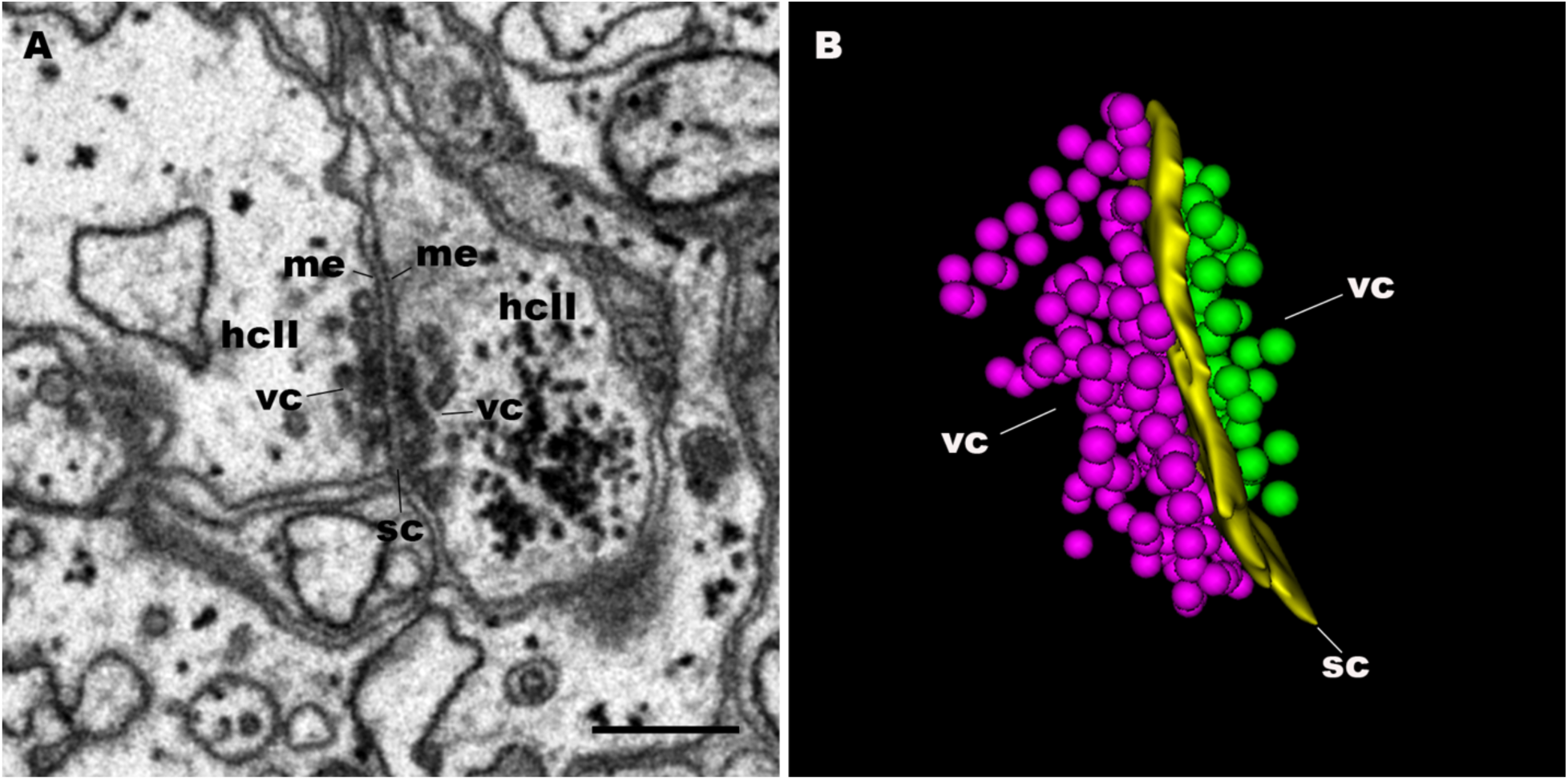
Structure of a two-way chemical synapse between type II hair cells. A serial block-face scanning electron microscopy (SBF-SEM) image (A) and a 3-D reconstruction (B) of a two-way chemical synapse between type II hair cells (hcII) shown in Figure 1M. Note in panel A that the synapse consists of two rigid, parallel membranes (me) sandwiching an approximately 30 nm wide, electron-dense synaptic cleft (sc). Panel B is a 3-D model of the synapse showing dense accumulation of synaptic vesicles, forming vesicle clouds (vc), on both sides of the synapse. Purple balls represent the vesicles of the type II hair cell on the left in panel A, and green balls represent those of the type II hair cell on the right. Scale bar: 500 nm

**Supplementary Figure 3:**
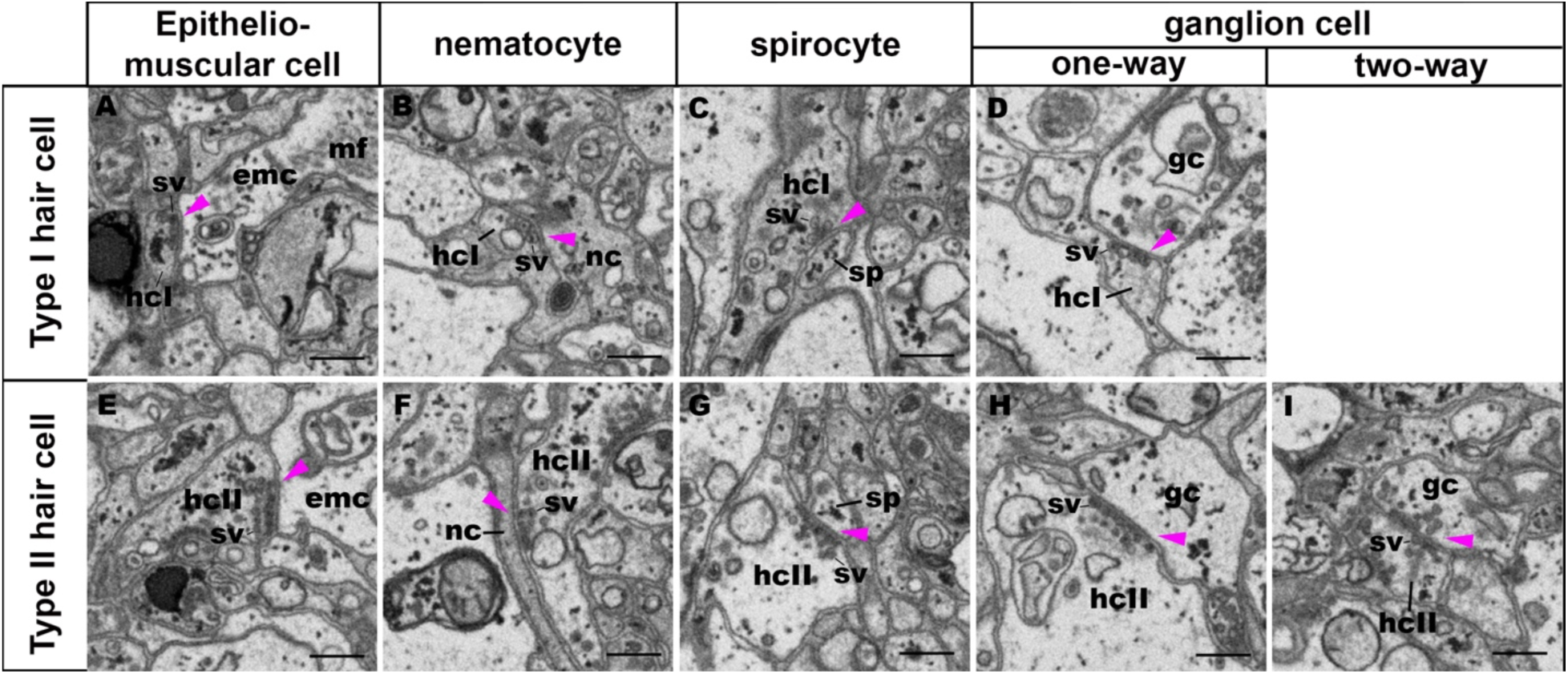
Synaptic connectivity of type I and type II hair cells. Serial block-face scanning electron microscopy (SBF-SEM) images of chemical synapses (purple arrowheads) of type I and type II hair cells (hcI and hcII, respectively). Both type I and type II hair cells are presynaptic to epitheliomuscular cell (emc; A, E), nematocyte (nc; B, F), spirocyte (sp; C, G), and ganglion cell (gc; D, H). In addition, type II hair cells, but not type I hair cells, form two-way synapses with ganglion cells (I). Note also that synaptic vesicles (sv) of type I hair cells are relatively large (ca. 95 nm) and primarily dense-cored (A-D), while those of type II hair cells are relatively small (ca. 70 nm), primarily opaque (E, G-I) and sometimes dense-cored (F). Scale bar: 500 nm

**Supplementary Figure 4:**
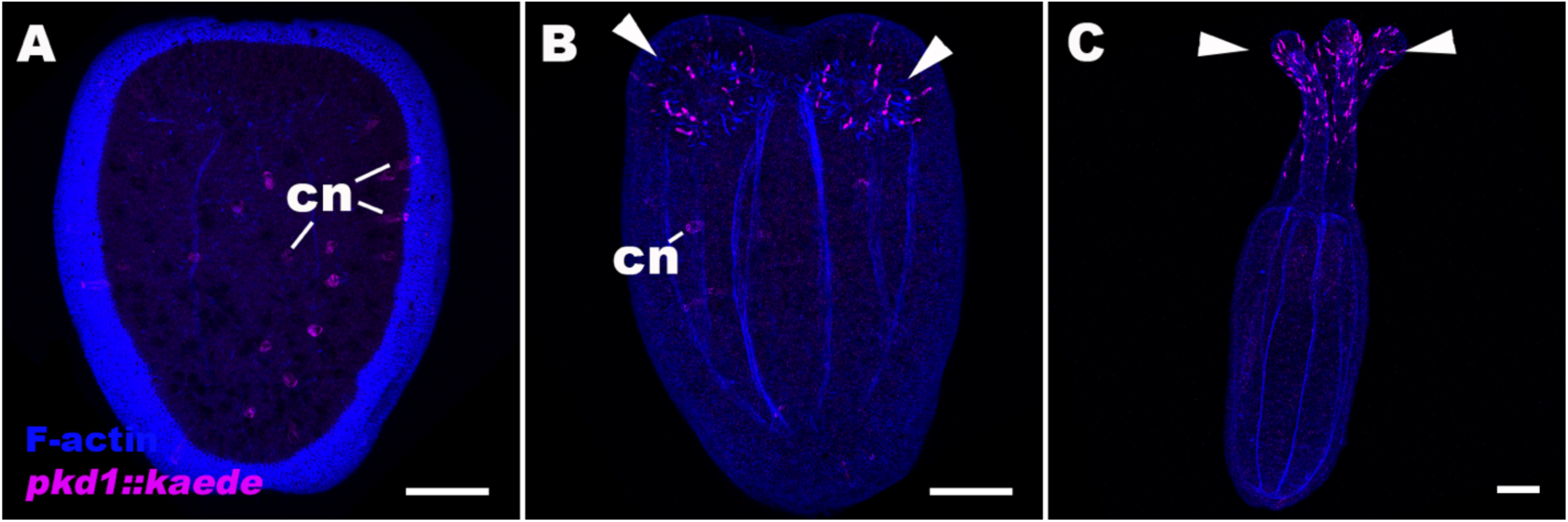
Developmental expression pattern of Kaede fluorescent protein in *polycystin-1::kaede* transgenic *Nematostella vectensis*. Confocal sections of *polycystin-1(pkd1)::kaede N. vectensis* at planula (A), tentacle-bud (B), and primary polyp (C) stages. Filamentous actin (“F-actin”) is labeled with a fluorescent dye SiR-Actin. All panels show side views of animals with the blastopore/mouth facing up. Panel A shows sections through the center of the animal, and panel B and C show sections through the entire animal. Low to moderate levels of *pkd1::kaede* expression are often observed in a subset of cnidocytes (cn) in the ectoderm during planula development through the tentacle-bud stage. *pkd1::kaede* expression in cnidocytes largely disappears by the primary polyp stage, while it becomes strongly expressed in a subset of ectodermal epithelial cells in tentacular ectoderm at life cycle transition - at the tentacle-bud and primary polyp stages (arrowheads in B and C). We note that cnidocyte expression of the transgene does not reflect the expression of endogenous *polycystin-1*; *in situ* hybridization with an antisense riboprobe against *polycystin-1* mRNA does not show cnidocyte expression (e.g.^1^), and there is no evidence from published cnidocyte transcriptome data (e.g.^2^) that *polycystin-1* is expressed in cnidocytes in planula larvae or adults. Scale bar: 50 µm

**Supplementary Figure 5:**
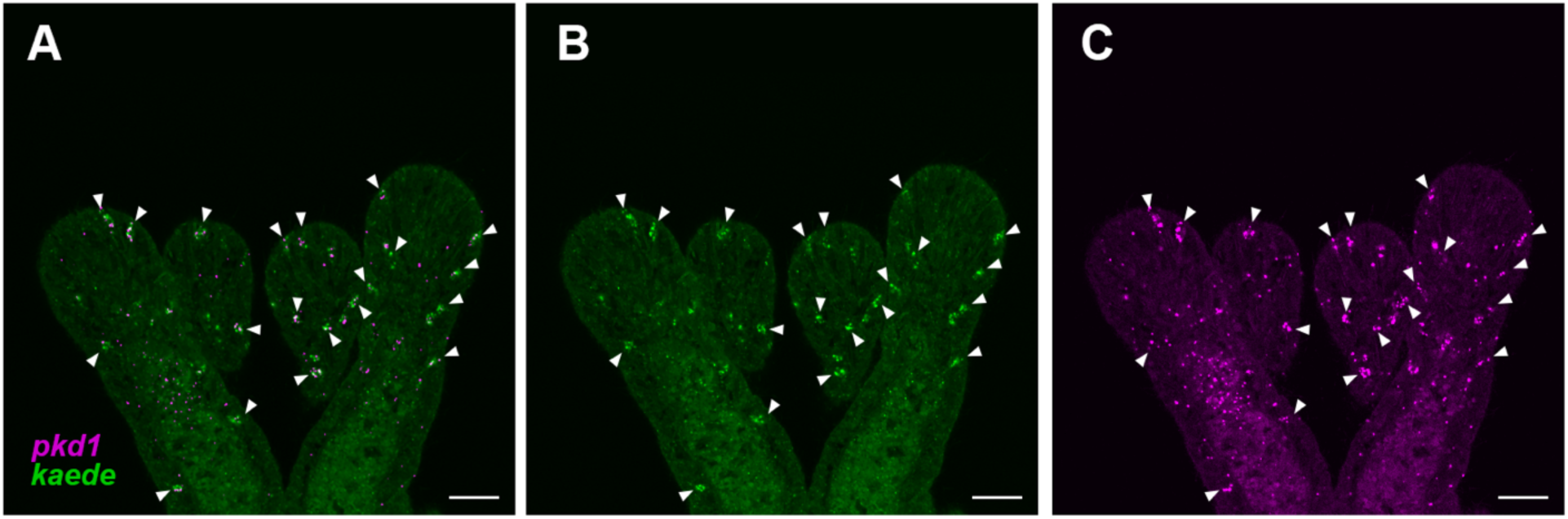
The expression pattern of *polycystin-1::kaede* recapitulates that of endogenous *polycystin-1* at the primary polyp stage. **A-C:** Confocal sections of a *polycystin-1::kaede* transgenic *N. vectensis* at the primary polyp stage, labeled with antisense riboprobes against *polycystin 1* transcript (‘pkd1’, shown in magenta) and *polycystin-1::kaede* transcript (‘kaede’ shown in green). Regions of overlapping signal are indicated with white arrows. Scale bar: 20 µm

**Supplementary Figure 6:**
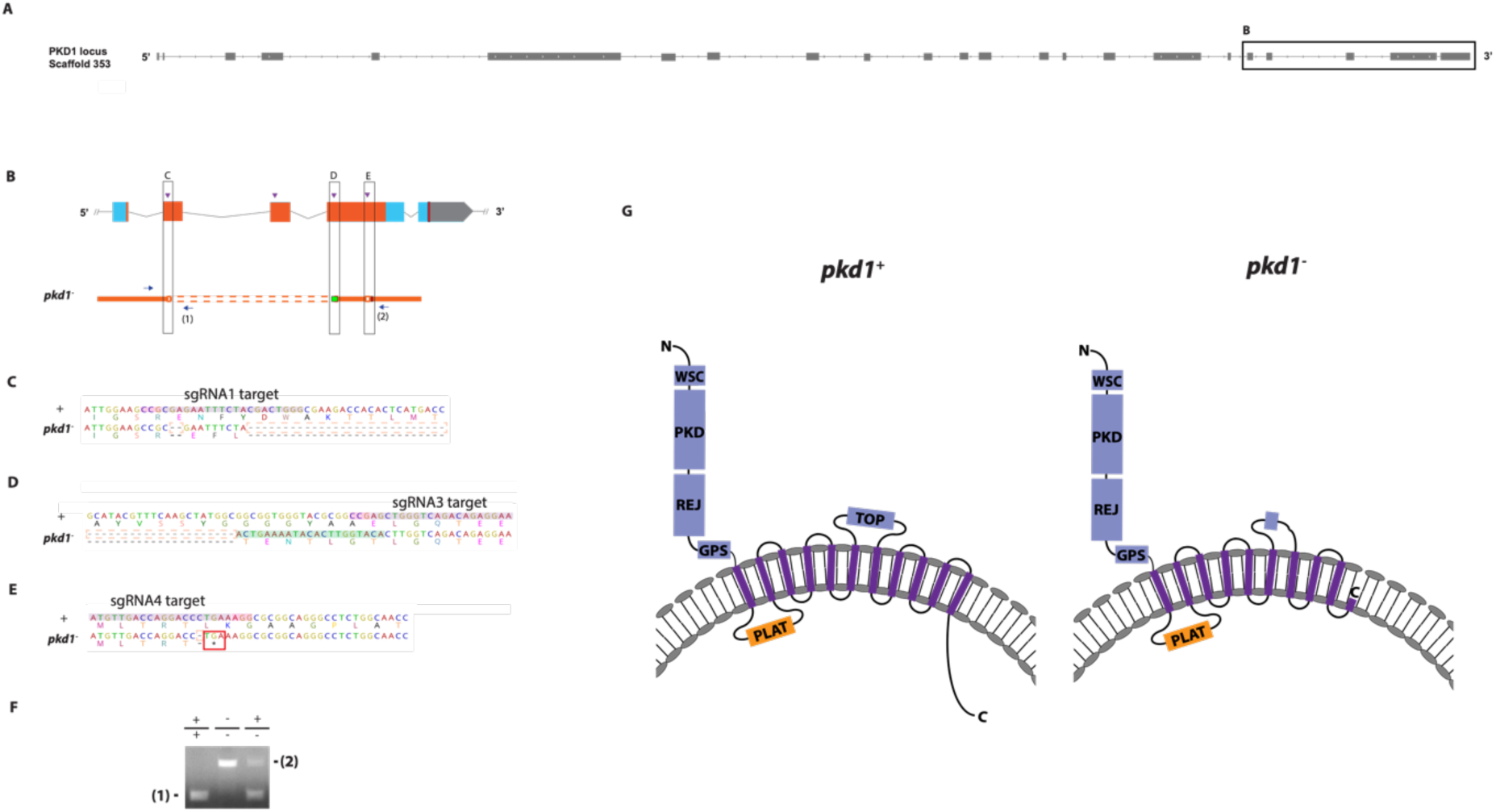
Generation of *polycystin-1* mutant sea anemone. A, B: Schematic views of the *polycystin-1/PKD1* locus and the mutant allele (*pkd1^−^*; B). Filled boxes represent 23 exons. Boxed region in A contains exons 19-23, expanded in B. In B, grey boxes are untranslated regions; aqua-colored boxes are translated regions. The region predicted to encode the Transient Receptor Potential (TRP) cation channel is highlighted in orange. Red bars show predicted translation termination sites, and purple arrowheads indicate sgRNA target sites. The *pkd1^−^*mutant allele contains a deletion mutation (dotted orange lines) and an insertion mutation (green). Blue arrows mark regions targeted in PCR analysis (F); reverse primers are numbered (1)-(2). C-E: nucleotide and translated amino acid sequences of wild-type and mutant alleles (from B). sgRNA target sites are shown in purple, PAM sites in pink, predicted translation termination site in red, insertions labeled in green, and deletions in dotted orange boxes. The mutant allele carries a 2 bp deletion mutation at the sgRNA1 target site, creating a translation frameshift (C), followed by a large deletion encompassing exons 20 and 22 (C, D), a 21 bp insertion mutation at the sgRNA3 target site (D), and a 1 bp deletion at the sgRNA4 target site generating a premature stop codon immediately after the mutation (E). F: Genotyping PCR. Primer (1) generates a 249 bp PCR product from the wildtype allele (’+’) but cannot bind to the *pkd1^−^* allele due to deletion mutation. Primer (2) generates a 829 bp PCR product from the *pkd1^−^*allele, and a 3554 bp PCR product from the wildtype allele. G: Schematics of wildtype (*pkd1+*) and mutant (*pkd1-*) Polycystin-1 protein. Extracellular domains are in blue, the intracellular domains in orange, and the transmembrane domains in purple. Domain prediction was performed as previously described^1^. The *pkd1-* mutant protein has a truncated TOP (tetragonal opening for polycystins) domain, and is missing the two C-terminal transmembrane domains thought to contribute to forming the channel pore^3^, and the C-terminal cytoplasmic tail. Abbreviations: WSC, cell wall integrity and stress response component; REJ, receptor for egg jelly; GPCR, G-protein coupled receptor; TM, transmembrane domain; PLAT, polycystin-1, lipoxygenase and alpha toxin

**Supplementary Figure 7:**
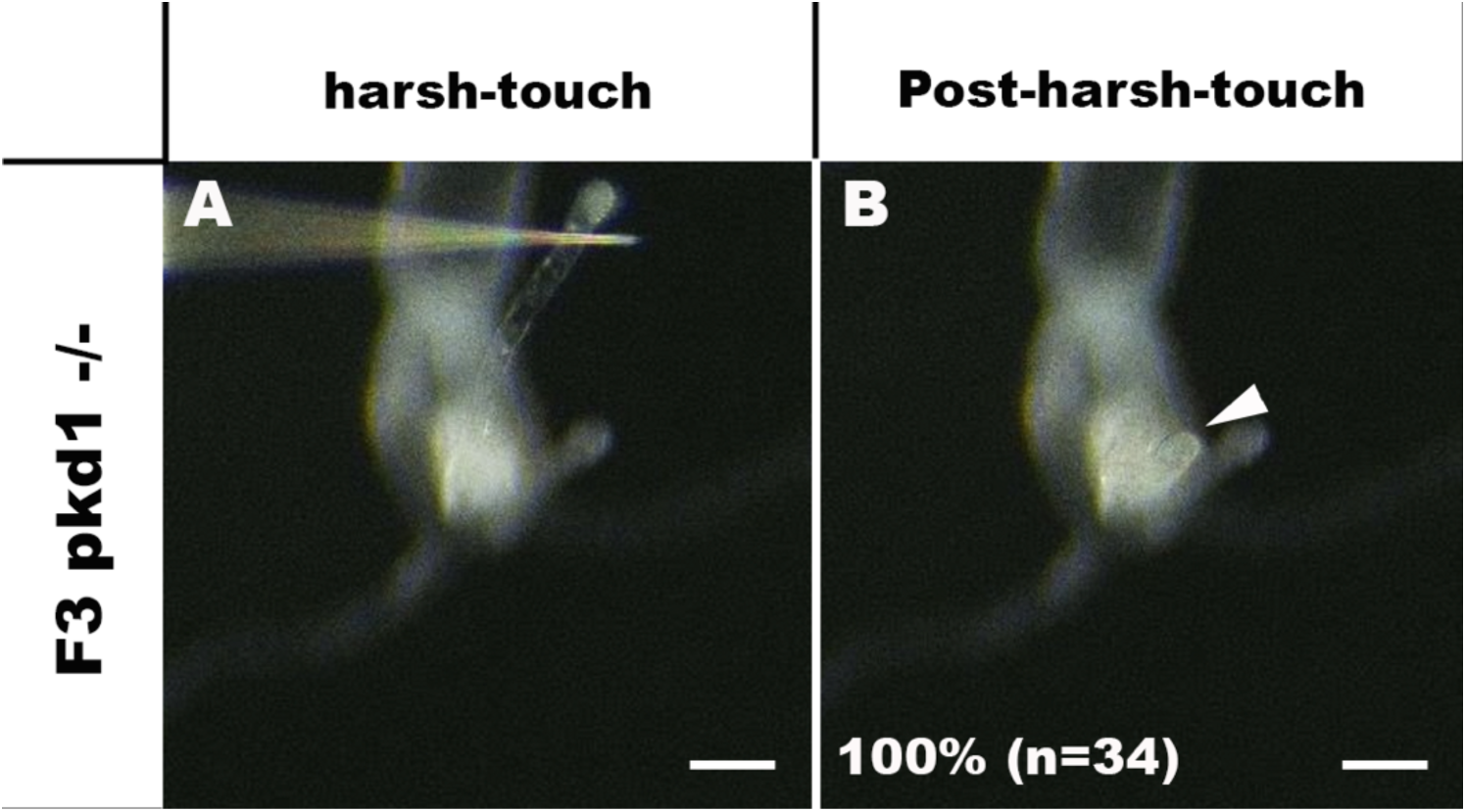
*polycystin-1* −/− homozygous mutants respond to harsh touch. Behavior of an F3 *pkd1* −/− *N. vectensis* polyp in response to harsh touch to its oral tentacle. Either a microinjection needle (shown) or a tungsten needle was used to press onto the surface epithelium of the tentacle to elicit harsh touch response – contraction of the stimulated tentacle. An animal before (A) and after (B) harsh touch are shown. All of the *pkd1* homozygous mutants tested were harsh touch-sensitive (100%, n=34), indicating that *polycystin-1* is not essential for harsh touch response. The arrowhead in B points to a stimulated tentacle exhibiting normal harsh touch response. Scale bar: 100 µm

**Supplementary Figure 8:**
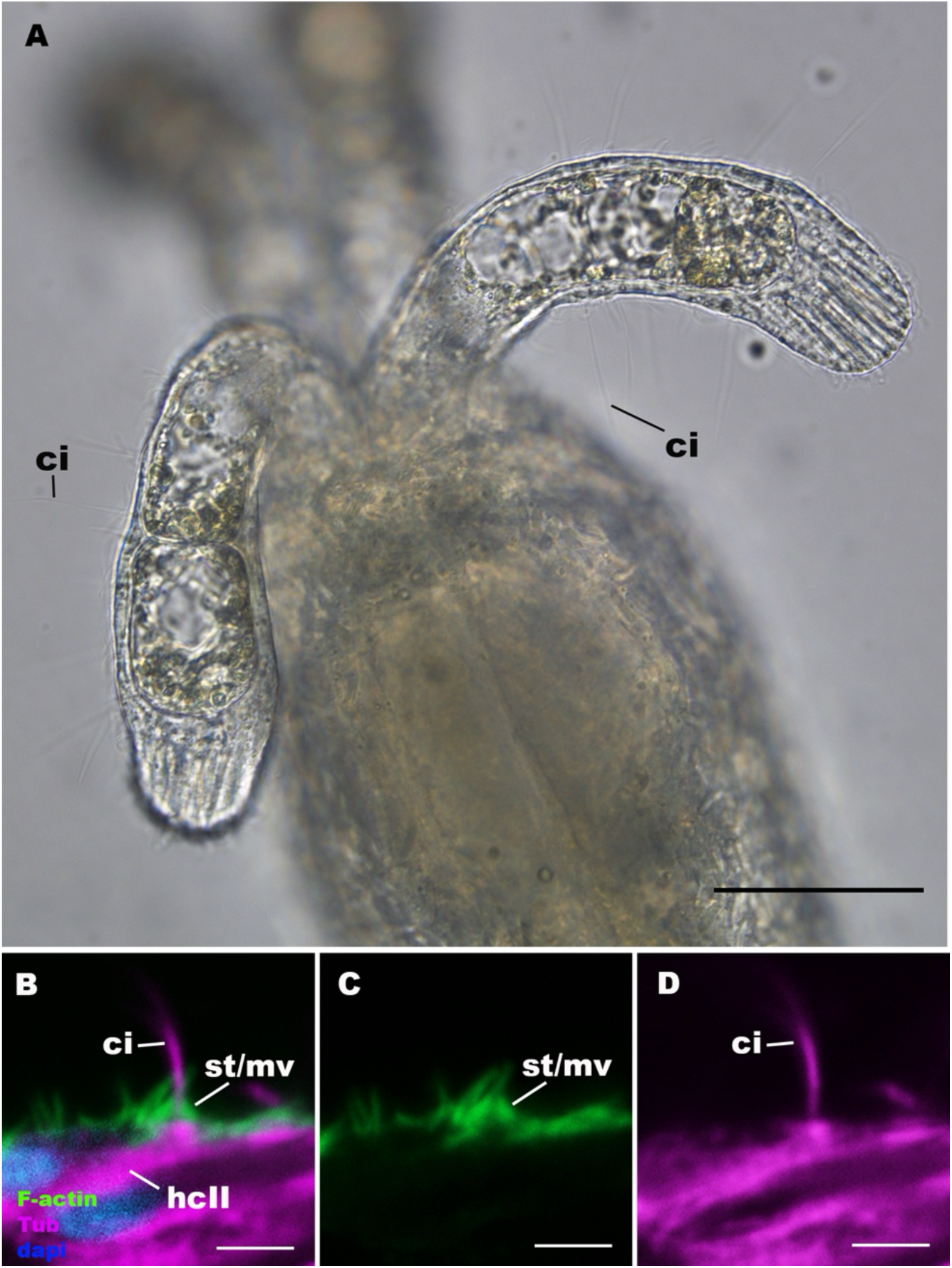
The apical sensory apparatus of type II hair cell develops in *polycystin-1* −/− homozygous mutant polyps. A: An optical section of oral tentacles of a *polycystin-1* −/− homozygous mutant polyp. The oral side is up. 20-40 µm long, erect cilia (ci) characteristic of type II hair cells emanate from tentacular ectoderm. B-D: Confocal sections of a type II hair cell (hcII) in a *polycystin-1* −/− homozygous mutant polyp, stained with antibodies against acetylated ∂-tubulin and tyrosinated ∂-tubulin (“Tub”). Filamentous actin is labelled with phalloidin (“F-actin”), and nuclei are labeled with DAPI (“dapi”). The apical epithelial surface faces up. Note the presence of an apical sensory apparatus consisting of a single cilium (ci) surrounded by rings of stereovilli/microvilli (st/mv). Scale bar: 50 µm (A); 5 µm (B-D

**Supplementary Figure 9:**
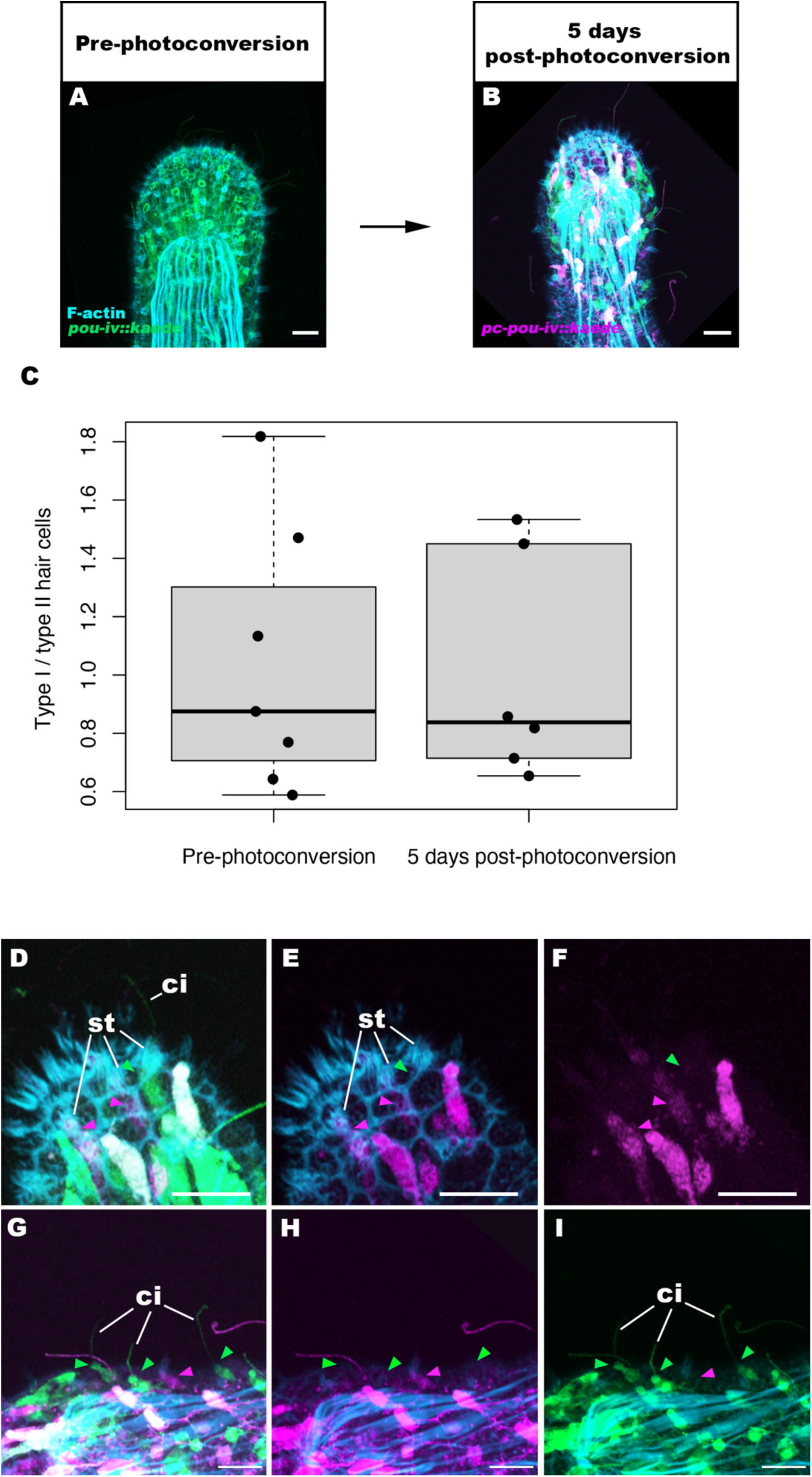
Cell tracking of type I and type II hair cells at the primary polyp stage provides no evidence of transformation between type I and type II hair cells. A, B: confocal sections of tentacles of *pou-iv::kaede* transgenic *N. vectensis* polyp. At the 16 dpf primary polyp stage (“Pre-photoconversion”; A), Kaede fluorescent protein (“*pou-iv::kaede*”) was photoconverted (“*pc-pou-iv::kaede*”) by ultraviolet illumination, and photoconverted kaede was used as a cell marker to examine the fate of *pou-iv::kaede*-positive type I and type II hair cells after 5 days (“5 days post-photoconversion”; B). Filamentous actin (“F-actin”) is labelled with a fluorescent dye SiR-Actin. Panels A and B show sections through the surface ectoderm; the distal end of the tentacle faces up. C: a box plot showing the proportions of type I hair cells relative to type II hair cells immediately before photoconversion (“Pre-photoconversion”; n=7 animals; mean 1.04) and 5 days after photoconversion (“5 days post-photoconversion”; n=6 animals; mean 1.00). Median (middle line), 25^th^, 75^th^ percentile (box) and 5^th^ and 95^th^ percentile (whiskers) are shown. Note that the proportion of type I hair cells relative to type II hair cells pre-photoconversion was determined based on the number of (non-photoconverted) *pou-iv::kaede-*positive cells, which is assumed to correspond directly to the proportion immediately after photoconversion (i.e. day 0), while the relative proportion of type I and type II hair cells 5 days post-photoconversion was calculated based on the number of photoconverted *pou-iv::kaede-*positive cells. The relative proportion did not significantly change (two tailed t-test, p=0.876), providing no support for the hypothesis that type I and type II hair cells represent different developmental states of the same cell type. D-I: magnified views of the tentacle shown in B. Note the presence of non-photoconverted type I (green arrowhead in D-F) and type II (green arrowheads in G-I) hair cells, characterized by pronounced stereovilli (st) and long cilia (ci), respectively; this indicates that the 5 day incubation period was sufficient to generate new type I and type II hair cells. Purple arrowheads show photoconverted type I hair cells. Scale bar: 10 µm

**Supplementary Figure 10:**
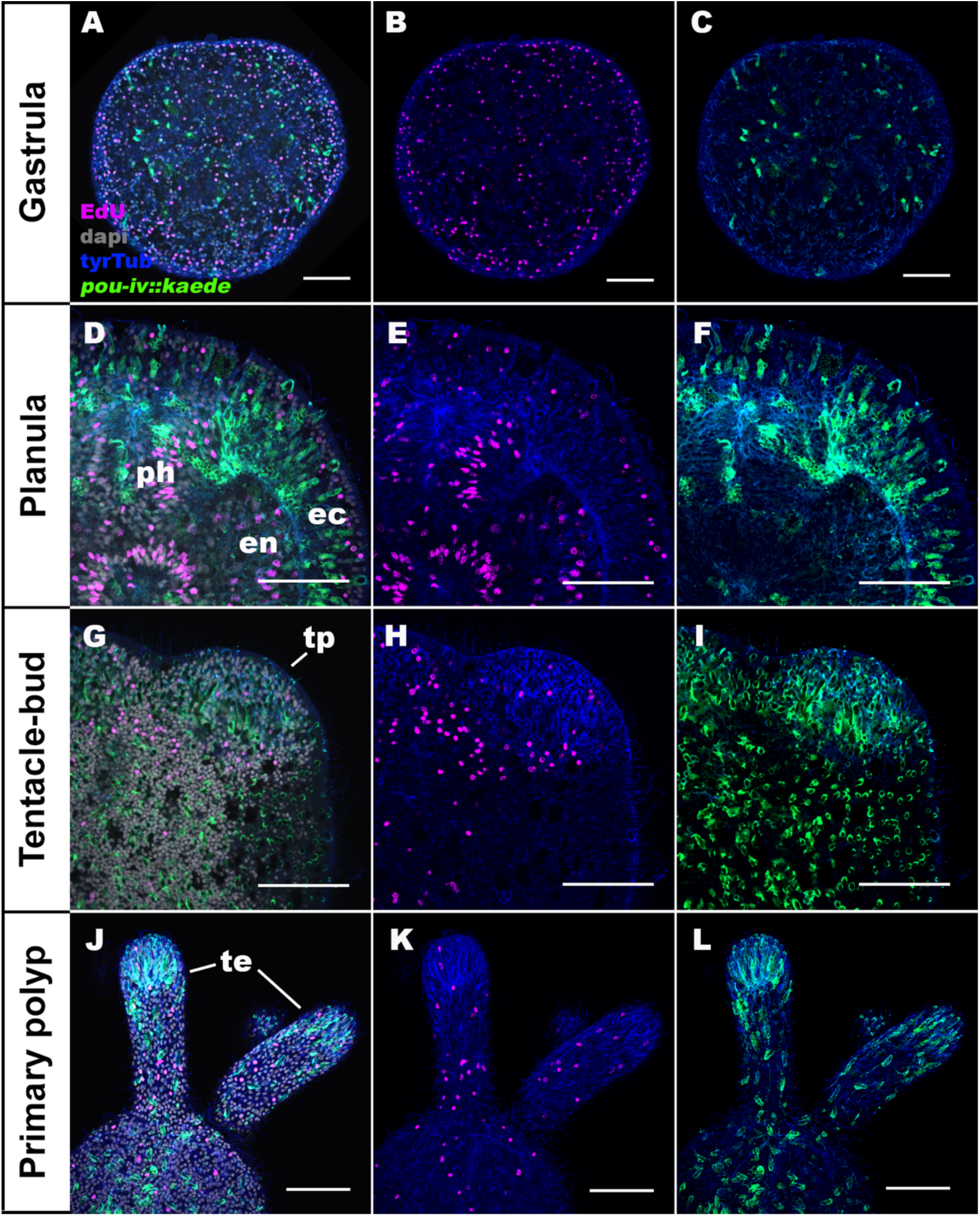
*pou-iv::kaede*-positive cells in the ectoderm are postmitotic. Confocal sections of *pou-iv::kaede* transgenic *N. vectensis* at gastrula (A-C), planula (D-F), tentacle-bud (G-I), and primary polyp (J-L) stages, labeled with antibodies against Kaede (*pou-iv::kaede*) and tyrosinated ∂-tubulin (“tyrTub”). S-phase nuclei are labeled by the thymidine analog EdU. In all panels, the blastopore/mouth faces up. A-C show sections through surface ectoderm of the entire embryo. D-F show medial sections of a circumoral region of the animal. G-I show sections of a tentacle primordium (tp) at the level of surface ectoderm. J-L show sections of oral tentacles (te) at the level of surface ectoderm. Consistent with postmitotic expression of endogenous *pou-iv* mRNA and protein reported previously^1,4^, *pou-iv::kaede-* positive cells in the ectoderm were EdU-negative across all developmental stages examined (gastrula n=2; early planula n=3; mid-planula I, n=3; mid-planula II, n=3; late planula, n=2; tentacle-bud, n=3; primary polyp, n=3; at least 30 *pou-iv::kaede-*positive cells were examined per animal for a total of 915 cells across stages), evidencing their postmitotic cell-cycle status. Abbreviations: ec ectoderm; en endoderm; ph pharynx. Scale bar: 50 µm

**Supplementary Figure 11:**
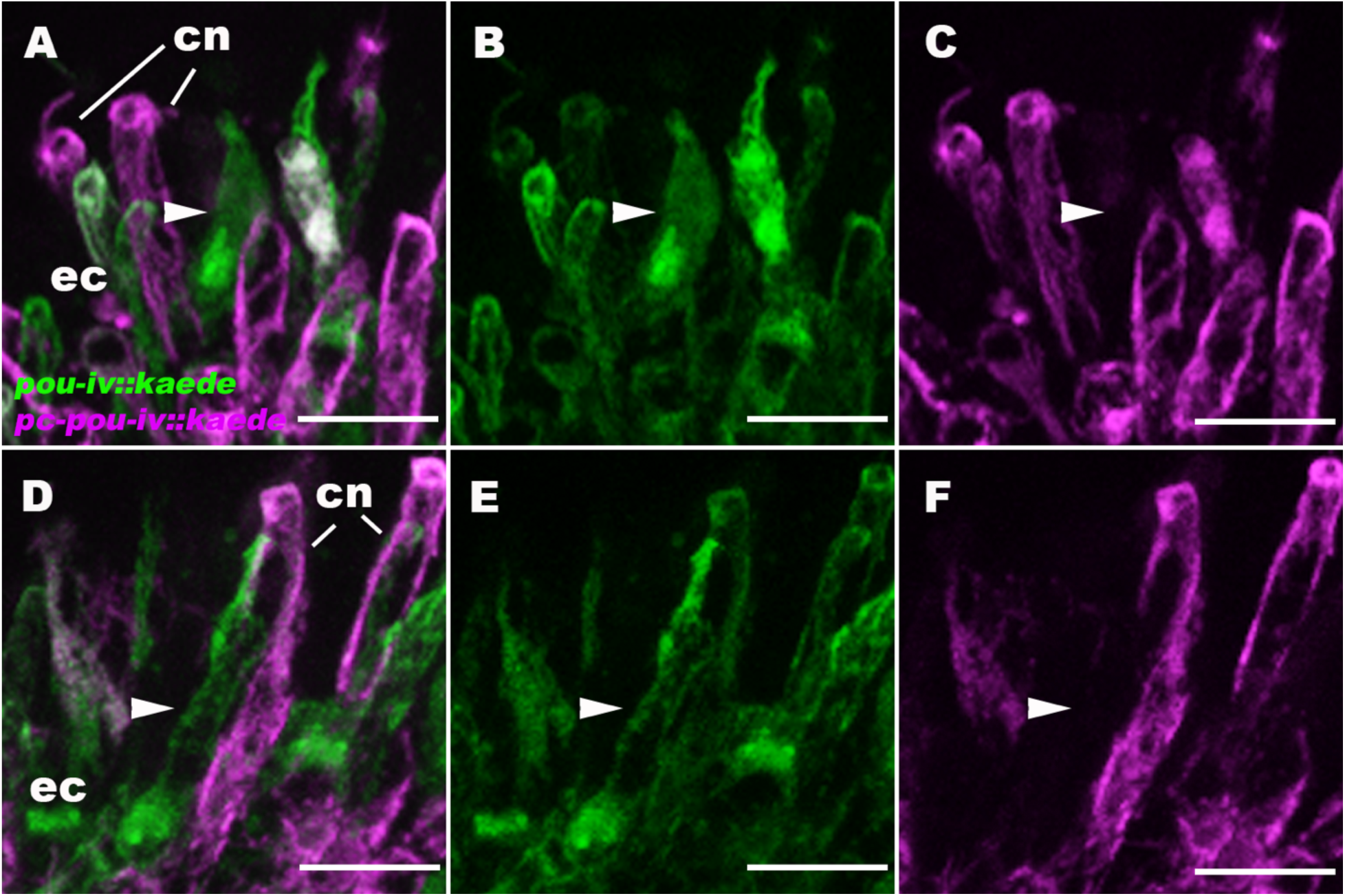
*pou-iv::kaede*-positive cells include morphologically immature cells. Confocal sections of tentacular ectoderm (ec) of *pou-iv::kaede* transgenic *N. vectensis* at the primary polyp stage. *pou-iv::kaede-*positive cells of the primary polyp were photoconverted at 12 dpf, and non-photoconverted Kaede (“*pou-iv::kaede”*) and photoconverted Kaede (“*pc-pou-iv::kaede”*) were imaged at 17 dpf. The apical epithelial surface faces up. While photoconverted cells are largely at morphologically mature states (e.g. cnidocytes (cn)), a subset of non-photoconverted Kaede-positive cells - which began expressing Kaede post-photoconversion – show immature morphology (arrowheads), evidencing that *pou-iv::kaede-*positive cells consist of postmitotic cells at different stages of maturation. Immature cells (arrowheads) are subepidermal in position, which suggests that tentacular sensory cells and cnidocytes establish apical contact with the external environment during maturation via basal-to-apical cell migration, cell shape changes or a combination of the two. Scale bar: 10 µm

**Supplementary Figure 12:**
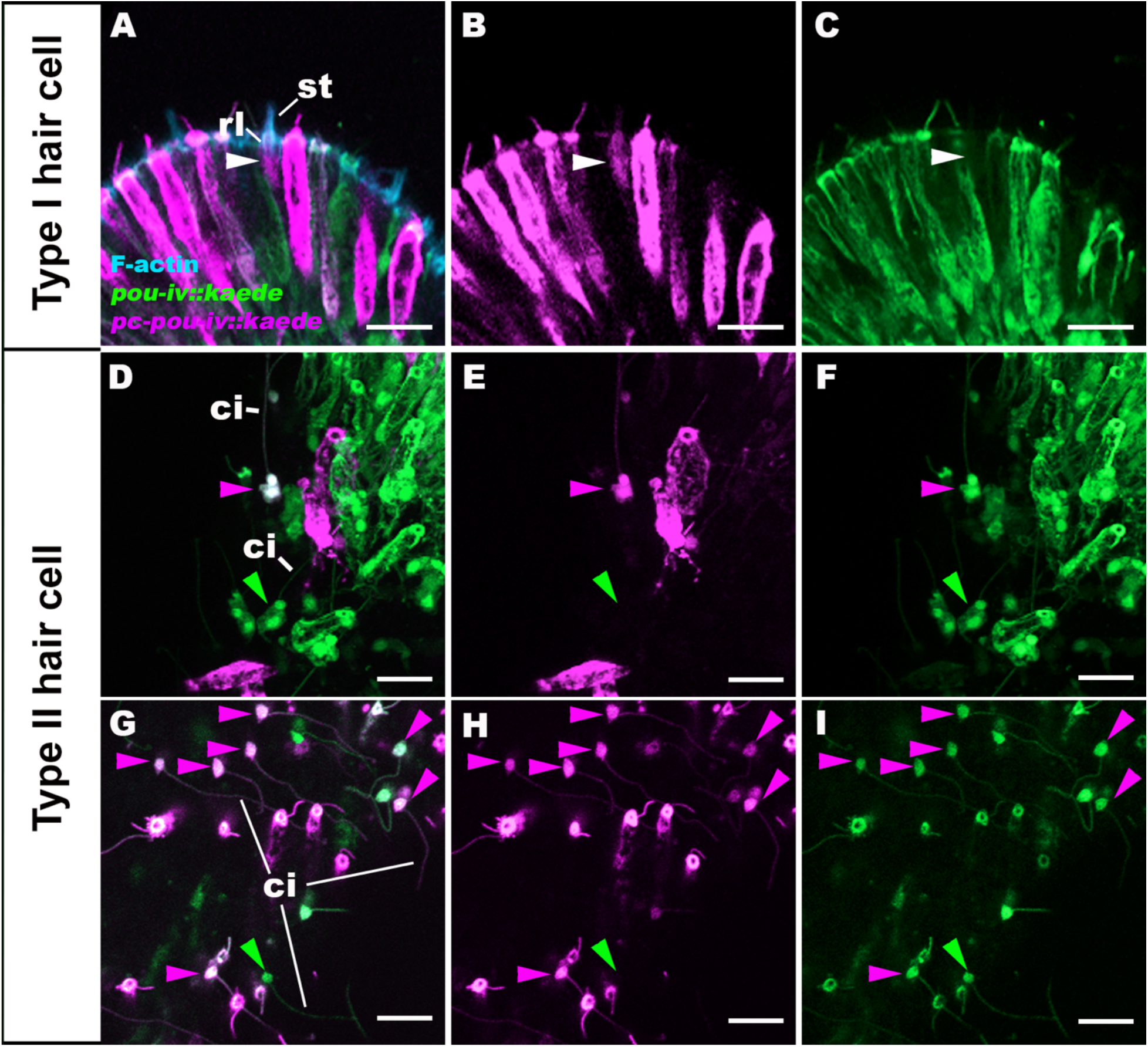
*pou-iv* promoter activity is differentially regulated in type I and type II hair cells. Confocal sections of tentacular ectoderm (ec) of *pou-iv::kaede* transgenic *N. vectensis* at the primary polyp stage. *pou-iv::kaede-*positive cells were photoconverted at the late planula (A-F) and primary polyp (G-I) stages, and non-photoconverted Kaede (“*pou-iv::kaede”*) and photoconverted Kaede (“*pc-pou-iv::kaede”*) were imaged at 5 days post-photoconversion. Filamentous actin (“F-actin”) is labeled with a fluorescent dye SiR-Actin. In A-C, the apical epithelial surface faces up. Panels D-I show superficial planes of section at the level of surface epithelium. The white arrowhead in A-C shows a photoconverted type I hair cell characterized by pronounced stereovilli (st) and their rootlets (rl). Note the lack of detectable levels of non-photoconverted Kaede in C, evidencing transient *pou-iv* promoter activities in type I hair cells. Purple arrowheads in D-I show photoconverted type II hair cells characterized by long cilia (ci); green arrowheads show non-photoconverted type II hair cells, which represent cells newly born after photoconversion. Note that photoconverted type II hair cells express non-photoconverted Kaede at levels comparable to newly born type II hair cells, evidencing persistent *pou-iv* promoter activities in type II hair cells. Scale bar: 10 µm

**Supplementary Figure 13:**
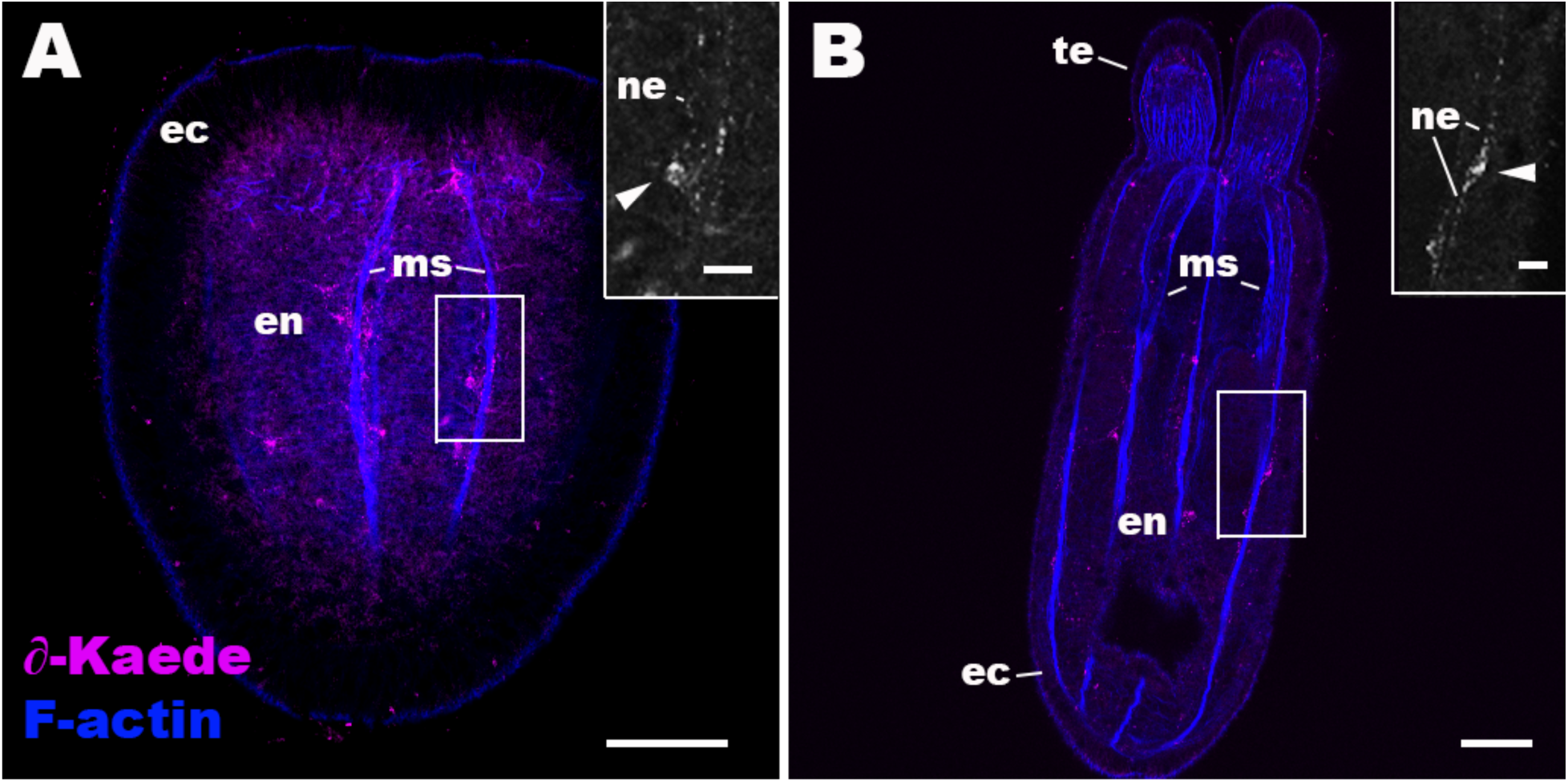
Anti-Kaede immunoreactivity occurs in a subset of endodermal neurons in wildtype *Nematostella vectensis*. Confocal sections of wildtype *N. vectensis* at the planula (A) and primary polyp (B) stages, labeled with an antibody against Kaede fluorescent protein (“∂-Kaede”). Filamentous actin is labeled with phalloidin (“F-actin”). Oral side is up. Sections are through the body column at the level of longitudinal retractor and parietal muscle fibers (ms). Insets show magnified views of anti-Kaede-immunoreactive endodermal neurons (arrowheads) with neurites (ne). Anti-Kaede immunoreactivity was not detectable in circumoral ectoderm of planula or tentacular ectoderm of primary polyp. Abbreviations: ec ectoderm; en endoderm; te tentacle. Scale bar: 50 µm; 10 µm (inset)

## Notes

### Competing Interest Statement

The authors have declared no competing interest.

